# Elevator Mechanism of Alternating Access in the *Escherichia coli* Concentrative Nucleoside Transporter NupC

**DOI:** 10.1101/2023.02.03.527023

**Authors:** Lijie Sun, Simon G. Patching

## Abstract

Members of the concentrative nucleoside transporter (CNT) family of proteins mediate uptake of nucleosides into cells driven by a cation gradient, which then enter salvage pathways for nucleic acid synthesis. In humans, they also transport hydrophilic anticancer and antiviral nucleoside analogue drugs into cells and tissues where they exert their pharmacological effects. *Escherichia coli* CNT NupC (400 residues) is pyrimidine-specific and driven by a proton gradient. We have used computational, biochemical, and biophysical methods to characterize evolutionary relationships, conservation of residues, structural domains, transmembrane helices, and residues involved in nucleoside binding and/or transport activity in NupC compared with those of sodium-driven *Vibrio cholera*e CNT (vcCNT) and human CNTs (hCNT1−3). As in the crystal structure of vcCNT, NupC appears to contain eight transmembrane-spanning α-helices. Wild-type NupC and single-cysteine-containing mutants were tested for transport activity in energized *E. coli* whole cells and for binding of nucleosides in non-energized native inner membranes using novel cross-polarization magic-angle spinning solid-state nuclear magnetic resonance methods. Wild-type NupC had an apparent affinity of initial rate transport (*K*_m_^app^) for [^14^C]uridine of 22.2 ± 3.7 *μ*M and an apparent binding affinity (*K*_d_^app^) for [1′-^13^C]uridine of 1.8−2.6 mM. Mutant S142C retained transport and binding affinities similar to those of the wild type. Mutants G146C and E149C had no transport activity but retained varying degrees of partial binding activity with affinities decreasing in the following order: wild type > S142C > G146C > E149C. Results were explained with respect to a homology model of NupC based on the structure of vcCNT and a hypothetical elevator-type mechanism of alternating access membrane transport in NupC.

## 1. Introduction

The concentrative nucleoside transporter (CNT) family of proteins, found in both prokaryotic and eukaryotic organisms, catalyzes inward movement of nucleosides across cell membranes against their concentration gradient, driven by a cation gradient moving in the same direction. In lower organisms, such as bacteria, CNTs scavenge nucleosides from the environment, and these enter salvage pathways for nucleic acid synthesis.^1^ In higher organisms, including humans, CNTs control cellular concentrations of physiological nucleosides; they also provide routes of entry for hydrophilic antiviral (e.g., zidovudine and ribavirin) and anticancer (e.g., Ara-C and gemcitabine) nucleoside analogue drugs.^2–7^ Nucleoside transporters are therefore important pharmacological targets, including sites in the brain.^8^ While this paper presents new methods and results with *Escherichia coli* proton-driven CNT NupC (400 residues),^9–13^ designs of experiments and discussions of results were also influenced by human CNTs and by other bacterial CNTs with available high-resolution crystal structures. Human CNTs are classified as solute carrier family SLC28;^14–18^ its three members (hCNT1−3) are sodium-driven and have selective substrate specificity and tissue distribution.^19–21^ hCNT1 and hCNT2 transport pyrimidine nucleosides (e.g., uridine, thymidine, and cytidine) and purine nucleosides (e.g., adenosine, guanosine, and inosine), respectively, coupling sodium:nucleoside transport with 1:1 stoichiometry. hCNT3 has broad selectivity for purine and pyrimidine nucleosides, coupling sodium:nucleoside transport with 2:1 stoichiometry. hCNT3 can also couple transport of uridine to a proton gradient with 1:1 stoichiometry.^22,23^

The first structurally characterized CNT was from the bacterium *Vibrio cholerae*.^24^ Sodium-driven vcCNT (418 residues) shares a high degree of sequence homology with hCNTs, the highest being 38.8% identity with hCNT3. The 2.4 Å crystal structure of vcCNT revealed a homotrimeric structural organization with each protomer comprising eight transmembrane-spanning α-helices (TM1−TM8), three interfacial helices (IH1−IH3) that run parallel to the membrane, and two hairpin re-entrant loops (HP1 and HP2) (Figure 1). Similar hairpin re-entrant loops were also seen in structures of the *Pyrococcus horikoshii* sodium-coupled aspartate transporter, GltPh, which also has eight transmembrane helices and exists as a homotrimer.^25–27^ The overall shape of the trimer is like an inverted triangular basin with its opening facing the intracellular side and a knoblike structure facing the extracellular side.^24^ The N- and C-terminal ends are both periplasmic, and each protomer has a novel overall fold divided into two subdomains based on locations at outer and inner regions relative to its center. The “scaffold domain” (TM1, TM2, IH1, EH, TM3, and TM6), in the outer region, maintains the overall architecture, including trimerization contacts. TM6 is exceptionally long (42 residues, ~60 Å) and tilted almost 60° with respect to the membrane normal. The “transport domain” comprises two structural groups (IH2, HP1, TM4, and TM5 and IH3, HP2, TM7, and TM8) on either side of TM6 that are related by internal two-fold pseudosymmetry. The tips of HP1 and HP2 and unwound regions of TM4 and TM7 lie at the center of the transport domain, located slightly below the middle of the membrane plane. These regions contain residues involved in direct binding interactions with nucleosides. Based on crystal structures of vcCNT in complex with uridine,^24,28^ HP1 residues Gly153, Gln154, Thr155, and Glu156 and TM4 residue Val188 interact with the base moiety, while HP2 residue Glu332 and TM7 residues Phe366, Asn368, and Ser371 interact with ribose. The single sodium-binding site (Asn149, Val152, Ser183, and Ile184) is located between the tip of HP1 and the unwound region of TM4, which is near the nucleoside-binding site.^24^ Crystal structures of a sodium-coupled CNT from *Neisseria wadsworthii* (nwCNT) (425 residues) have been determined in outward-facing, multiple intermediate, and inward-facing conformations.^29^ nwCNT shares 66.5% sequence identity with vcCNT, and its structures have confirmed that CNTs appear to adopt an elevator-type mechanism of alternating access membrane transport,^30–36^ first hypothesized from crystal structures of GltPh.^25,26,37^

**Figure 1.**
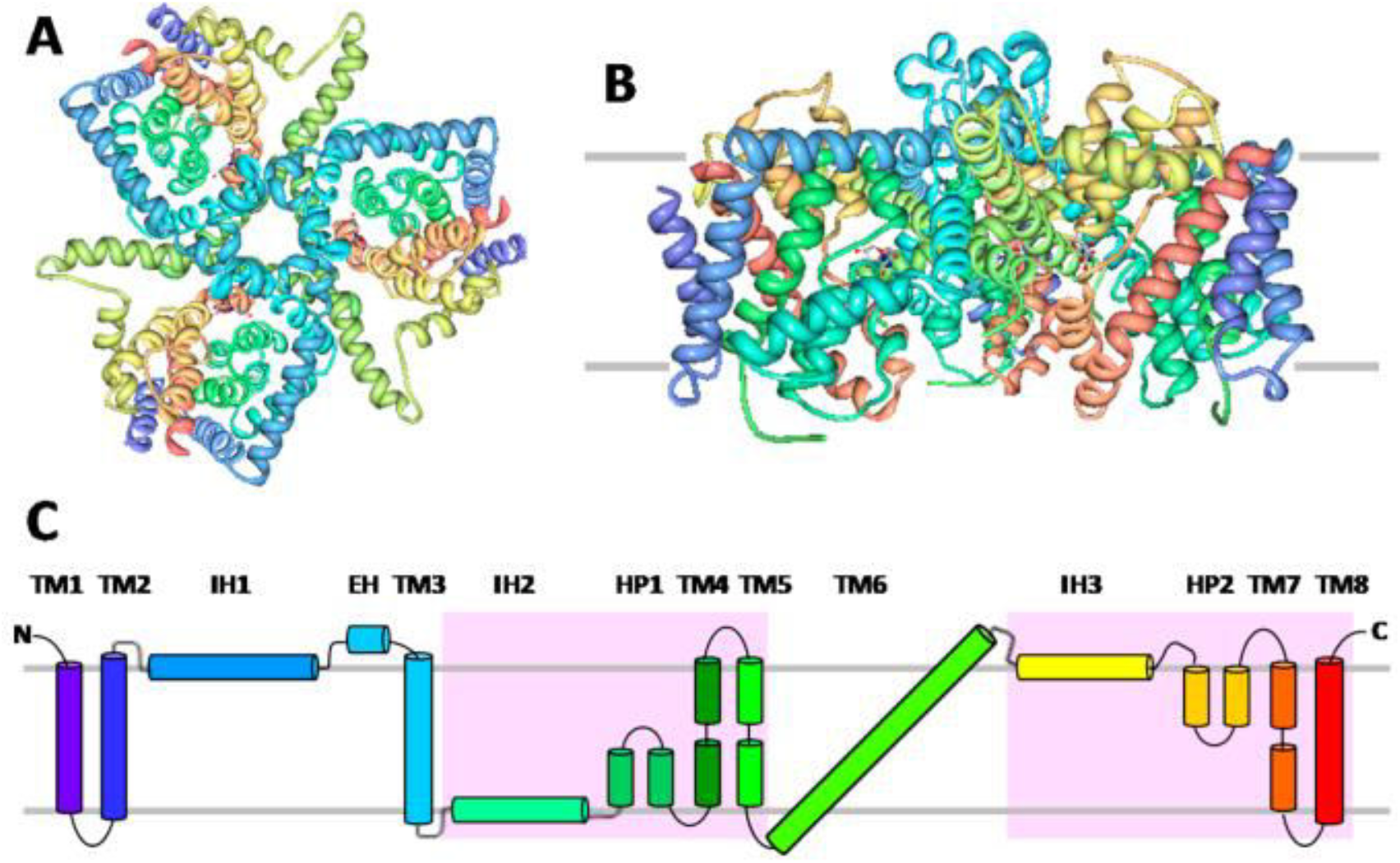
Structural organization of the *V. cholerae* concentrative nucleoside transporter vcCNT. Crystal structure of homotrimeric Na^+^-coupled nucleoside transporter vcCNT from *V. cholerae* in complex with uridine viewed (A) from the periplasm and (B) from the membrane plane. In panel B, the grey horizontal lines indicate the boundaries of the membrane. The structure was drawn with the N-terminus coloured purple and the C-terminus coloured red using Protein Data Bank entry 4PD6 and PDB Protein Workshop 3.9.^53^ (C) Membrane topology of vcCNT. Schematic illustration of transmembrane-spanning helices (TM1−TM8), interfacial helices (IH1−IH3), extracellular helices (EH), and hairpin re-entrant loops (HP1 and HP2) in vcCNT based on the crystal structure. The “transport domain” is highlighted in pink.

NupC transports the pyrimidines uridine, cytidine, and thymidine and also the purine adenosine.^9–13^ While NupC does not transport guanosine and inosine, there is evidence to suggest transport of xanthosine with a very low affinity.^38^ Heterologous expression of NupC in *Xenopus laevis* oocytes demonstrated a nucleoside selectivity resembling that of human and rat CNT1, including an ability to transport adenosine and the therapeutic drugs 3′-azido-3′-deoxythymidine (AZT, zidovudine), 2′,3′-dideoxycytidine (ddC, zalcitabine), and 2′-deoxy-2′,2′-difluorocytidine (gemcitabine).^39^ Apparent affinities for transport (*K*_m_^app^ values) were 1.5−6.3 *μ*M for adenosine, uridine, and gemcitabine and 112 and 130 *μ*M for AZT and ddC, respectively. Competitive inhibition of [^14^C]-uridine transport by recombinant NupC in *E. coli* whole cells with 46 unlabelled physiological nucleosides and nucleobases, nucleoside analogues, and other compounds facilitated rigorous investigation of ligand recognition.^40^ This assay identified specific structural motifs necessary for binding of ligands by NupC compared with *E. coli* major facilitator superfamily nucleoside transporter NupG.^41–44^ The overall pattern of inhibition demonstrated the close functional relationship of NupC with hCNTs and confirmed NupC as a useful experimental model for the human proteins.^40^ Using nucleosides synthesized with nuclear magnetic resonance (NMR)-active isotope labels (^13^C and ^15^N),^40,45^ solid-state NMR methods were developed for observing and characterizing direct binding of nucleosides to NupC with amplified expression in native *E. coli* inner membranes. Under these non-energized conditions, the apparent *K*_d_ value for binding of [1′-^13^C]uridine to NupC was 2.6 mM.^46^

This work explores evolutionary relationships, sequence homologies, and structural and functional characteristics of NupC compared with those of vcCNT, nwCNT, and hCNTs. This includes analyses of NupC residues putatively involved in direct binding interactions with nucleosides and the predicted membrane topology and structural organization of NupC based on the structure of vcCNT. Site-specific mutants of NupC were constructed and characterized, including measurement of transport activities and nucleoside binding affinities using novel solid-state NMR methods. Results are explained with respect to a homology model of NupC and a hypothetical elevator-type mechanism of alternating access concentrative nucleoside transport.

## 2. Materials and Methods

### 2.1. Computational Methods

Protein sequences were taken from the UniProt KnowledgeBase (http://www.uniprot.org/) and aligned using Clustal Omega (http://www.ebi.ac.uk/Tools/msa/clustalo/).^47^ Phylogenetic analysis was performed by exporting the resultant neighbour-joining phylogenetic tree in Newick format and redrawing using iTOL: Interactive Tree of Life (http://itol.embl.de/).^48^ Membrane topology predictions were performed using TOPCONS (http://topcons.cbr.su.se/pred/),^49^ TMHMM (http://www.cbs.dtu.dk/services/TMHMM/),^50^ and CCTOP (http://cctop.enzim.ttk.mta.hu/).^51^ Amino acid compositions were determined using the ExPASy tool ProtParam (http://web.expasy.org/protparam/).^52^ Protein structures were drawn using PDB Protein Workshop 3.9 and NGL viewer.^53,54^ The homology model of NupC was produced using ExPASy tool SWISS-MODEL (https://swissmodel.expasy.org/).^55^

### 2.2. Site-Directed Mutagenesis

Site-directed mutagenesis used the QuikChange polymerase chain reaction (PCR) method (Stratagene). “Touchdown” PCR was typically performed over 18−20 cycles using the following conditions: denaturation at 94 °C for 15 s, annealing at 60−55 °C for 30 s, and extension at 72 °C for 30 s per kilobase of template DNA followed by final extension at 72 °C for 10 min. An aliquot (10 *μ*L) of the reaction mixture was analyzed by agarose gel electrophoresis, and a further aliquot (20 *μ*L) was treated with 10 units of *Dpn*I at 37 °C for 1−2 h to degrade parental methylated (5′-Gm^6^ ATC-3′)and hemimethylated DNA. An aliquot (1−2 *μ*L) of the *Dpn*I-treated reaction mixture was used to transform competent *E. coli* Top10 cells (50 *μ*L). These were transferred to LB-agar plates containing carbenicillin (100 *μ*g/mL) and incubated overnight at 37 °C. Colonies containing putative mutant plasmids were grown overnight in LB medium (5 mL) and DNA minipreps performed. The presence of the required mutation was confirmed by DNA sequencing. Cysteineless NupC (C96A) was constructed first and used as the parental protein for construction of all further mutants.

### 2.3. Cell Growth

*E. coli* strain BL21(DE3) harboring isopropyl *β*-D-1-thiogalactopyranoside (IPTG)-inducible plasmid pGJL16 for amplifying expression of wild-type (WT) NupC or constructs with the necessary mutations for producing NupC mutants (C96A, C96A/S142C, C96A/G146C, or C96A/E149C) were grown in M9 minimal medium supplemented with glucose (20 mM) in batch culture (500 mL volumes in 2 L baffled flasks) at 37 °C while being shaken at 200 rpm. At the optimal stage of growth established in pilot studies, IPTG (0.5 mM) was added to induce amplified expression. Typically, cells were grown to an *A*_600_ of ~0.6 over 4 h, followed by a further period of 3 h for induction to a final *A*_600_ of ~1.8−2.0.

### 2.4. Transport Studies on Whole Cells

Measurements of the rate of uptake of [^14^C]uridine (Amersham Biosciences) into energized *E. coli* whole cells expressing WT NupC or mutant forms were performed using the method described previously.^40^ Data were obtained during initial rates of [^14^C]uridine transport after an uptake period of 15 s, and the resultant curves were used to estimate values of *K*_m_ and *V*_max_ by fitting to the Michaelis−Menten equation using GraphPad Prism. For expression tests, aliquots of the same cells used for transport measurements were subjected to mixed membrane preparation by the water lysis method^56,57^ and analysis by Western blotting.

### 2.5. Western Blotting

Samples of mixed membranes containing 10 *μ*g of total protein were resolved by sodium dodecyl sulfate-polyacrylamide gel electrophoresis (SDS−PAGE), and proteins were detected by Western blotting using a primary antibody raised against a 15-residue sequence of NupC (SEENIQMSNLHEGQS).^58^ Proteins were transferred from the gel to a nitrocellulose membrane by blotting at a constant voltage (100 V) for 1 h using a Bio-Rad Mini Protean II apparatus and a buffer containing glycine (190 mM), Tris (25 mM), and methanol (20%). The blot was washed twice for 10 min in Tris-buffered saline {TBS [20 mM Tris-HCl and 500 mM NaCl (pH 7.5)]} and then twice for 20 min in TTBS (TBS containing 0.1% Tween 20). The blot was blocked by being washed with non-fat dried milk (5%) in TTBS for 1 h and then washed twice for 10 min in TTBS. The blot was incubated overnight with a 1:1000 dilution of the primary antibody in antibody buffer (1% non-fat dried milk in TTBS) and then washed three-times for 15 min in TTBS. The blot was incubated for 1 h with a 1:5000 dilution of the secondary antibody (anti-rabbit IgG horseradish peroxidise-conjugated antibody, Sigma) in antibody buffer and then washed three-times for 15 min in TTBS. The blot was removed from the wash, drained, and then transferred to a sheet of Saran wrap on a flat surface. A solution of chemiluminescent substrate (Supersignal West Pico chemiluminescent substrate, Pierce) (~5 mL) was transferred to the blot to cover it evenly and then left at room temperature for 5 min. The blot was drained of solution, enveloped in Saran wrap, and exposed to X-ray film for 1 min.

Removal of any contaminating antibodies from the primary antibody preparation was achieved by treatment with an acetone powder of *E. coli* total protein before use. An aliquot (5 *μ*L) of a primary antibody solution was diluted to 1 mL with TBS, and 10 mg [1% (w/v)] of a total protein acetone powder from *E. coli* BL21(DE3) cells was added. The mixture was incubated at 4 °C on a blood-wheel for 30 min and then spun in a microcentrifuge (10000 rpm, 10 min) to remove the protein. The supernatant containing the primary antibody was diluted to the required titer and then used to probe the Western blot as described above.

### 2.6. Membrane Preparation

Inside-out vesicles were prepared from harvested cells by explosive decompression in a French press, and the inner membrane fraction was isolated by sucrose density gradient ultracentrifugation.^57^ Membrane vesicles were washed three-times by suspension in 20 mM Tris-HCl buffer (pH 7.5) followed by ultracentrifugation before final suspension in the same buffer for storage at −80 °C after rapid freezing. The total protein concentration in the vesicle suspension was determined by the method of Schaffner and Weissmann, and the percent NupC content was estimated by densitometry measurements on the proteins resolved by SDS−PAGE and stained with Coomassie Brilliant Blue. Control membranes without amplification of NupC expression were prepared in the same way from cells containing plasmid pTTQ18 without the transporter gene insert.

### 2.7. NMR Conditions

^13^C-observed cross-polarization magic-angle spinning (CP-MAS) NMR measurements were preliminarily performed at the University of Leeds using a Varian InfinityPlus 300 instrument operating at a magnetic field of 7 T with a Varian double-bearing MAS probe tuned to 75.32 MHz for ^13^C and 299.5 MHz for ^1^H. Measurements were repeated and continued at the University of Manchester Institute of Science and Technology (UMIST) (which was subsumed by the University of Manchester) and at the University of Liverpool using a Bruker Avance 400 spectrometer operating at a magnetic field of 9.3 T, corresponding to Larmor frequencies of 400.13 MHz for ^1^H and 100.13 MHz for ^13^C. All experiments used a sample temperature of 4 °C. At 400 MHz, standard ^1^H−^13^C CP-MAS experiments were performed using a double-tuned probe head at a sample spinning frequency of 4 kHz. CP was initiated by a ^1^H 90° pulse length of 3.5−4.0 *μ*s, followed by Hartmann−Hahn cross-polarization from ^1^H to ^13^C at a ^1^H field of 65 kHz. Continuous wave proton decoupling at a field of 85 kHz was applied during signal acquisition. Variable contact time experiments used contact times of 0.1, 0.5, 1, 2, 3, 5, 8, and 10 ms. Spectra were acquired over the given number of scans; chemical shifts were referenced to adamantane at 37.8 ppm, and spectra were processed with 40 Hz exponential line broadening.

Spectra representing [1′-^13^C]uridine interacting specifically with NupC were obtained by difference after displacement of the bound substrate with either 40 or 80 mM unlabelled uridine. Spectra were first obtained at the different contact times from membranes containing 5 mM [1′-^13^C]uridine. Unlabelled uridine was then added to the membranes, and another series of spectra was obtained under identical NMR conditions. In the second series of spectra, the [1′-^13^C]uridine was fully displaced from NupC but a residual peak from [1′-^13^C]uridine was present, which was assigned as labelled substrate interacting non-specifically with the membranes. Each of the second series of spectra was subtracted from the corresponding spectrum obtained at the same contact time before unlabelled uridine was added. This produced a series of spectra exhibiting a peak from [1′-^13^C]uridine representing only the fraction of the labelled substrate interacting with NupC. These difference spectra are termed “displacement spectra”.

### 2.8. Analysis of NMR Data

Values of the dissociation constant (*K*_d_) representing the affinity of [1′-^13^C]uridine for NupC were estimated from two independent NMR experiments. In one experiment, peak intensities for [1′-^13^C]uridine were measured from ^13^C CP-MAS NMR spectra of membranes containing 0−80 mM unlabelled uridine at a fixed contact time. Intensities were plotted as a function of unlabelled uridine concentration and apparent *K*_d_ values calculated using a standard Marquardt−Levenberg curve fitting procedure. The second experiment used variable contact time CP-MAS measurements, whereby peak intensities for specifically bound [1′-^13^C]uridine were measured from displacement spectra obtained with a range of contact times. Values of the apparent *K*_d_ and the rate constant for substrate dissociation (*k*_off_) were estimated by comparing experimental intensities with simulations obtained using the Monte Carlo method.^46^ Simulated curves were a function of *k*_off_ and *K*_d_, contact time (*t*_c_), substrate and protein concentrations, ^1^H *T*_1_ relaxation times in the rotating frame (*T*_1*ρ*_) for the free and bound substrate, and the CP rate constant for the bound substrate (*R*_HC_). A series of curves was calculated for contact times of 0−10 ms at different combinations of *k*_off_ and *K*_d_ values, keeping all the other input parameters constant. Values of *T*_1*ρ*_ and *R*_HC_ were measured experimentally or estimated from natural abundance signals of other membrane components as described previously.^46^ Curves in closest agreement with experimental data were determined by minimizing the χ^2^ function.

## 3. Results and Discussion

### 3.1. Evolutionary Relationships and Conservation of Residues in NupC and CNT Family Proteins

The close evolutionary relationship between NupC and other characterized CNT family proteins (vcCNT, nwCNT, and hCNTs) is demonstrated by a nearest-neighbor phylogenetic tree annotated with levels of sequence homology (Figure S1). NupC is most closely related to vcCNT and nwCNT (30.3 and 32.3% identical and 28.8 and 27.3% highly similar, respectively; combined 59.1 and 59.6%, respectively) and then with the human proteins in the following order: hCNT3 > hCNT1 > hCNT2. Based on separate sequence alignments, human CNTs share 35.9% identical residues and 25.0% highly similar residues (combined 60.9%) (Figure S2), with the closest relationship between hCNT1 and hCNT2 (63.3% identical, 17.7% highly similar, combined 81.0%) (Table S1). Like NupC, vcCNT is most closely related to hCNT3 (38.8% identical, 25.4% highly similar, combined 64.2%) (Table S1). A BLAST search for NupC against all Eukaryota in the UniProt KnowledgeBase identified two very close homologues from the soil fungus *Beauveria bassiana* (Figure S3). Proteins A0A0A2W106 (395 residues) and A0A0A2V7L6 (380 residues) have sequences that are 92.7 and 73.4% identical to that of NupC, respectively. No other reliable close homologues of NupC were identified, with all other eukaryotic proteins having sequences less than 30% identical with that of NupC.

A sequence alignment between NupC and vcCNT highlights conserved residues, locations of vcCNT structural domains (TM1−TM8, IH1−IH3, HP1, and HP2) and vcCNT residues involved in direct binding interactions with nucleosides (Figure S4). The region of vcCNT structural domains involved in nucleoside binding (HP1, TM4, HP2, and TM7) is highly conserved with NupC and hCNTs (Figure S5). Of nine vcCNT residues involved in direct binding interactions with nucleosides, seven are identically conserved in NupC and two are highly similar (Table 1). Also note that all nine vcCNT residues are identically conserved in nwCNT. The corresponding residues in NupC are Gly146, Gln147, Ser148, Glu149, Ile181, Glu321, Phe350, Asn352, and Ser355. Except for hCNT1 residues corresponding to Gly153 and Val188 and the hCNT2 residue corresponding to Gln154, all vcCNT residues involved in nucleoside binding are identically conserved in hCNTs (Table 1). Of four vcCNT sodium-binding residues, three (Asn149, Val152, and Ile184) are identically conserved in hCNTs but not in NupC. The position corresponding to vcCNT sodium-binding Ser183 is occupied by threonine in NupC and hCNTs. These differences are consistent with NupC being proton-driven and vcCNT, nwCNT, and hCNTs being sodium-driven.

**Table 1.**
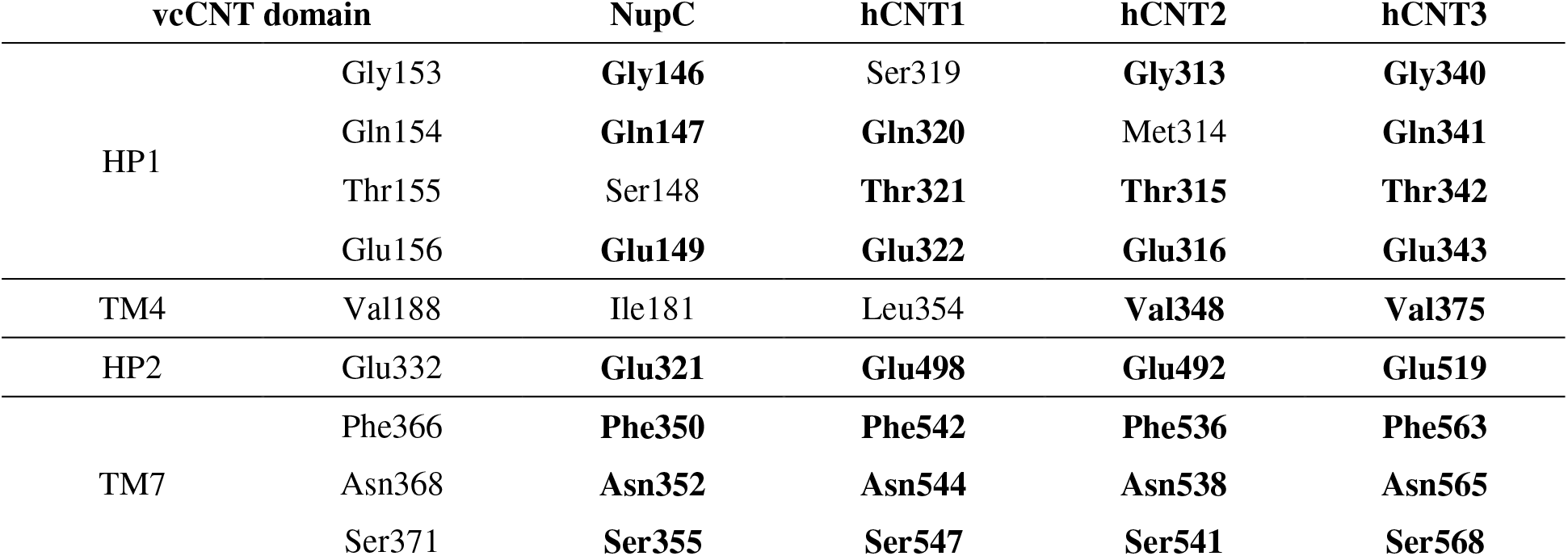
Conservation of Nucleoside-Binding Residues in Concentrative Nucleoside Transporters. Residues in *V. cholerae* vcCNT structural domains (HP1, TM4, HP2, and TM7) that interact directly with nucleosides based on crystal structures of vcCNT^24,28^ are compared with residues at the corresponding positions in NupC from *E. coli* (P0AFF2) and human concentrative nucleoside transporters hCNT1 (O00337), hCNT2 (O43868), and hCNT3 (Q9HAS3) based on a sequence alignment (Figure S3). Residues that are identically conserved are shown in bold and those that are highly similar in italic. Binding site residues in vcCNT are all identical with those in nwCNT.

Residues responsible for nucleoside specificities in hCNT1 and hCNT2 are those corresponding to Ser319/Gln320 and Ser353/Leu354 in hCNT1. Conversion to the corresponding residues in hCNT2 (Gly313/Met314 and Thr347/Val348) changes hCNT1 from a pyrimidine-specific to a purine-specific transporter.^19^ Furthermore, hCNT1 mutation S353T produces a profound decrease in cytidine transport efficiency and, in combination with L354V (S353T/L354V), produces a novel uridine-preferring transport phenotype.^59^ On this basis, similar mutations at corresponding positions in NupC are predicted to change NupC from pyrimidine-specific to purine-specific. The position in NupC corresponding to hCNT1 residue Ser319 is already glycine (Gly146), and also glycine in hCNT3 (Gly340), which may explain the ability of NupC to transport adenosine. Other NupC mutations predicted to alter nucleoside specificity would be Q147M, S180T, and I181V. Roles for hCNT1 residues Glu308, Glu322, and Glu498 in sodium-coupled nucleoside transport have also been identified, with Glu322 having the strongest influence on hCNT1 transport function.^60^ Corresponding residues at all three of these positions are identically conserved as glutamate in NupC, vcCNT, hCNT2, and hCNT3 (Figure S6).

### 3.2. Predictions of Transmembrane Helices and Distributions of Amino Acids in NupC and vcCNT

To demonstrate challenges associated with defining the structural organization of NupC, its sequence was subjected to analysis by three of the most popular and/or best performing tools that predict transmembrane helices: TOPCONS,^49^ TMHMM,^50^ and CCTOP^51^ (Figure S7). TOPCONS has been updated to efficiently separate any N-terminal signal peptides from transmembrane regions and can predict re-entrant loops.^61^ Also, the ability of TOPCONS to predict 1551 transmembrane helices in 235 membrane proteins with high-resolution structures has recently been assessed, giving a 94.8% success rate.^62^ For NupC, the TOPCONS consensus result predicted nine transmembrane helices with a periplasmic N-terminus and a cytoplasmic C-terminus (Figure S7A). TMHMM predicted eight full transmembrane helices with periplasmic N- and C-termini and also a ninth partial helix (Figure S7B). The CCTOP consensus result predicted nine transmembrane helices with a cytoplasmic N-terminus and a periplasmic C-terminus (Figure S7C). The positions of the nine transmembrane helices predicted by these three methods are roughly in agreement with each other or at least show significant overlap (Figure S8). Arrangements predicted by TOPCONS and TMHMM are most plausible based on the “positive-inside rule”.^63^ Figure S8 also shows putative locations of transmembrane helices in NupC based on sequence homology and the structure of vcCNT. While transmembrane helices based on vcCNT all show some overlap with those coming from the prediction tools, there are significant differences in their lengths and in their start and end positions. This is partly because the prediction tools generally give the same number of residues in all transmembrane helices, for example, 22 and 21 residues from TOPCONS and TMHMM, respectively.^62^ The prediction tools were therefore never going to identify the full length of a 42-residue helix for TM6 as in vcCNT. A further limitation of the prediction tools is that they tend to miss transmembrane helices containing internal positively charged residues.^64^ The presence of positively charged residues may therefore have affected their ability to recognize locations for many of the putative transmembrane helices in NupC based on those provided by the structure of vcCNT (Figure S8). The extra “ninth” helix in NupC given by the prediction tools appears in the region of residues 278−307 (Figure S8), corresponding to IH3 in vcCNT (Figure S5). All three prediction tools gave a full transmembrane helix for NupC at this location. A high-resolution crystal structure of NupC is clearly desirable to confirm whether it shares the same structural organization as vcCNT.

For vcCNT, both TOPCONS and TMHMM predicted ten transmembrane helices with periplasmic N- and C-termini (Figure S9A,B). CCTOP predicted eight transmembrane helices with periplasmic N- and C-termini (Figure S9C). Of ten transmembrane helices predicted by TOPCONS and TMHMM, eight show good agreement or significant overlap, while the others show significant discrepancies. These eight helices in the TOPCONS and TMHMM results all show significant overlap with the eight transmembrane helices predicted by CCTOP (Figure S8). While the eight common helices coming from the prediction tools all show some overlap with transmembrane helices in the structure of vcCNT, they all have different lengths and different start and end positions. The differences may partly be due to the presence of internal positively charged residues in some of the transmembrane helices in vcCNT. The structure of vcCNT does conform to the “positive-inside rule”,^63^ with the majority of positively charged residues confined to the cytoplasmic side (Figure 2). Negatively charged residues have an approximately equal distribution at the periplasmic and cytoplasmic sides of vcCNT (Figure 2). In terms of other types of amino acids in vcCNT, hydrophobic residues predominate in transmembrane regions, as expected. Hydroxyl-containing residues and amido-containing residues are scattered throughout the transmembrane helices and the periplasmic and cytoplasmic regions (Figure 2).

**Figure 2.**
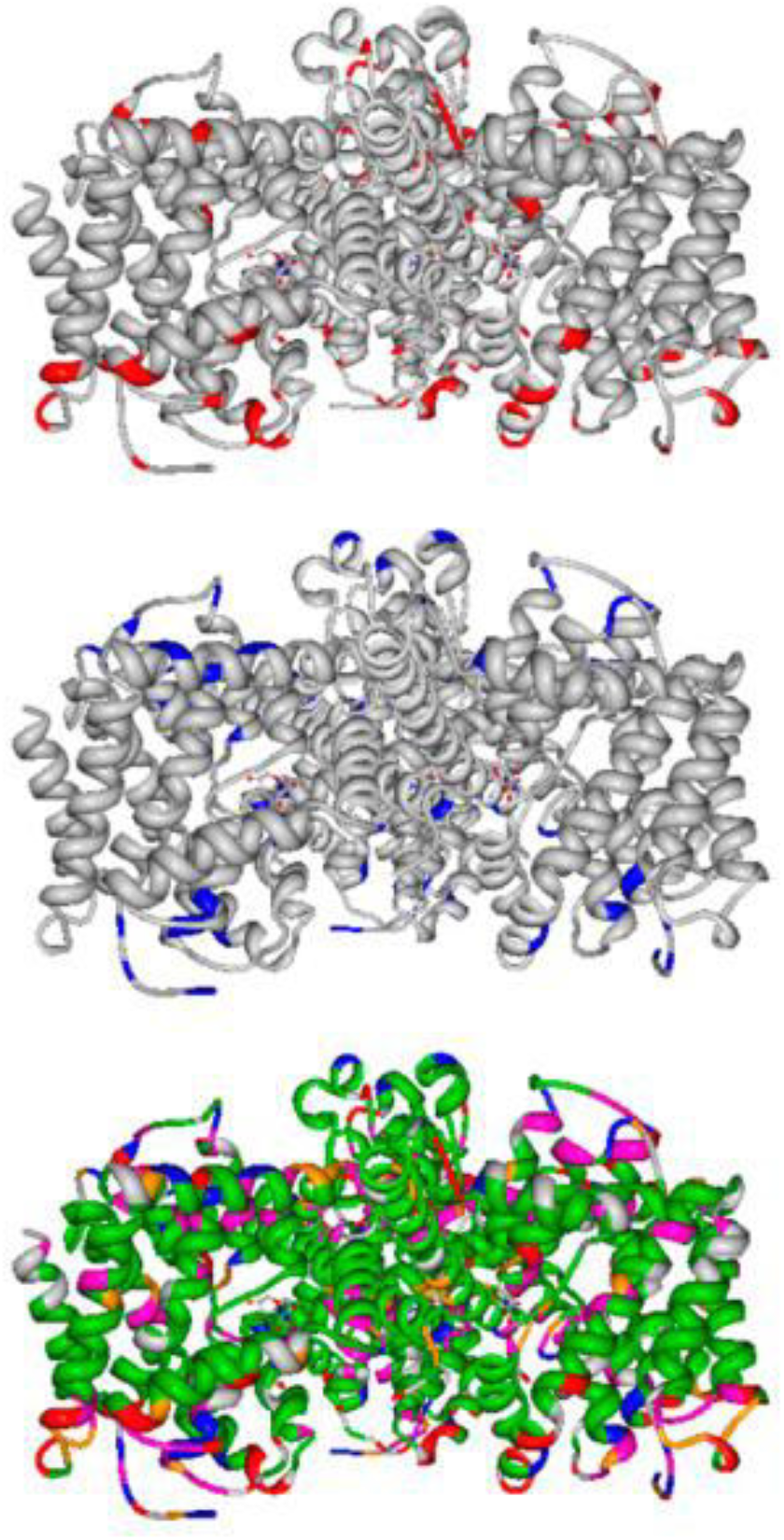
Distribution of amino acids in the structure of vcCNT from *V. cholerae*. The crystal structure of vcCNT from *V. cholerae* in complex with uridine was drawn using Protein Data Bank entry 4PD6 and PDB Protein Workshop 3.9.^53^ Residues are coloured to show those that are positively charged at physiological pH (arginine and lysine) (*red*), negatively charged at physiological pH (aspartic acid and glutamic acid) (*blue*), hydrophobic (alanine, isoleucine, leucine, phenylalanine, valine, and glycine) (*green*), hydroxyl-containing (serine, threonine, and tyrosine) (*pink*), and amido-containing (*orange*).

Amino acid compositions in NupC, vcCNT, nwCNT, and hCNTs were also calculated (Table S2). Percentage contents for individual amino acids and for groupings of amino acids with similar physicochemical properties (hydrophobic, positively charged, negatively charged, hydroxyl-containing, and amido-containing) in NupC, vcCNT, and nwCNT were similar to overall average values in 235 secondary transport proteins from *E. coli*.^65^ Percentage contents in hCNTs were similar to overall average values in 336 human secondary transport proteins.^64^ Hence, none of the CNTs contain significantly high contents of any individual type of amino acid or group of amino acids with similar physicochemical properties.

### 3.3. Characterization of NupC Cysteine Mutants at Positions Involved in Nucleoside Binding and Transport Activity

#### 3.3.1. Details and Construction of NupC Cysteine Mutants

Cysteine scanning mutagenesis investigated roles of specific NupC residues in nucleoside binding and transport activity.^66^ This work deals with three residues (Ser142, Gly146, and Glu149) in a region of NupC corresponding to HP1 in the structure of vcCNT (Figure 3). Ser142 corresponds to Asn149 in vcCNT (and nwCNT), which is involved in coordinating the sodium ion (Figure 3), and this position is conserved as asparagine in hCNTs (Figure S5). Gly146 is conserved in vcCNT, nwCNT, hCNT2, and hCNT3 and corresponds to Ser319 in hCNT1 (Figure S5) that when mutated to glycine allows purine nucleoside transport.^19^ The corresponding vcCNT residue (Gly153) is close to both the base moiety of uridine^28^ and the sodium-binding site (Figure 3). Glu149 corresponds to Glu156 in vcCNT, which coordinates with the base moiety of uridine via water molecules,^28^ and this position is conserved as glutamate in hCNTs (Figure S5). Interestingly, mutant E149D retained partial transport activity, suggesting an important role for a carboxyl group at this position.^66^ NupC contains one native and non-essential cysteine residue (Cys96) for which the corresponding position in vcCNT, nwCNT, and hCNTs is occupied by proline (Figure S6). This residue was mutated to alanine to create cysteineless NupC (C96A), which retained the same transport activity as WT,^66^ and this was used as the parental protein for construction of the cysteine mutants. Three single-cysteine-containing mutants (S142C, G146C, and E149C) were constructed, expressed in *E. coli* (Figure S10), and characterized for nucleoside transport activity, nucleoside binding, and the effect of thiol-specific reagents.

**Figure 3.**
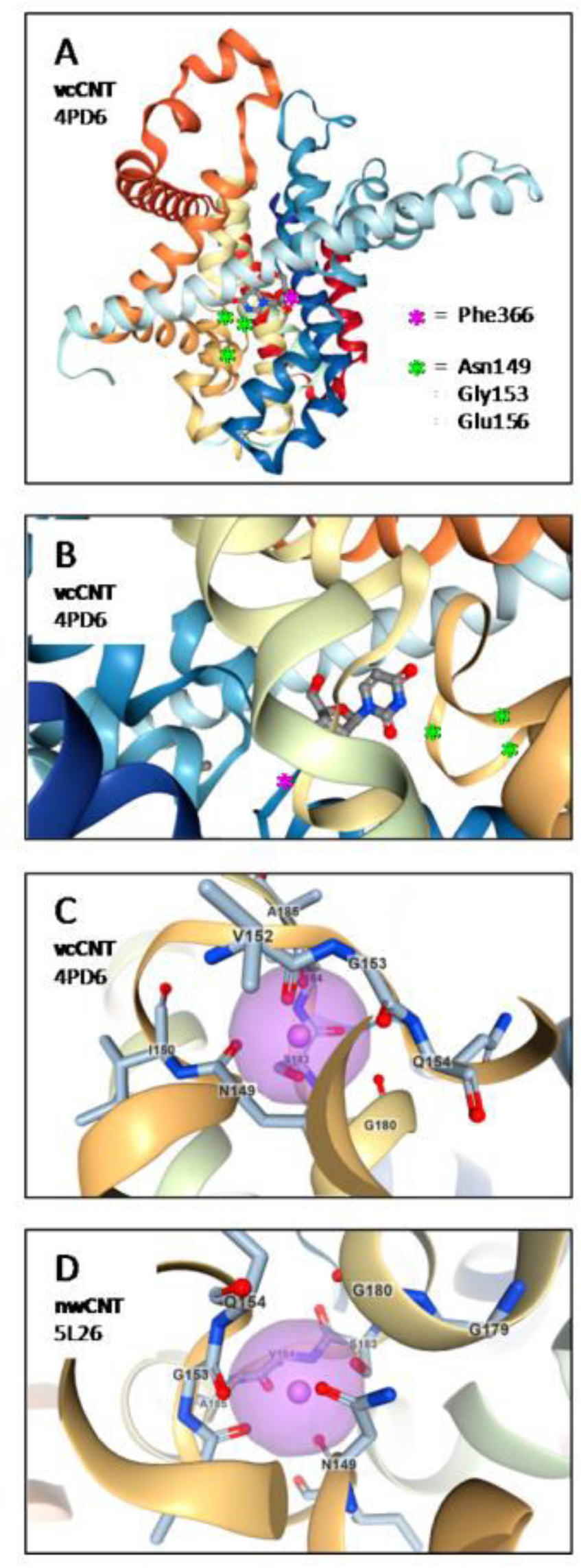
Locations of vcCNT and nwCNT residues corresponding to NupC mutated residues Ser142, Gly146, and Glu149. (A) Model of a vcCNT protomer with bound uridine indicating the positions of Asn149, Gly153, and Glu156 in HP1 that correspond to mutated NupC residues. The position of crucial TM7b residue Phe366 is also indicated. (B) Model of the vcCNT uridine-binding site showing positions of residues Asn149, Gly153, and Glu156 in relation to the base moiety of uridine. The position of important Phe366 is also indicated. (C) Model of the vcCNT sodium-binding site showing the positions of residues Asn149, Gly153, and Glu156. (D) Model of the nwCNT sodium-binding site showing the positions of residues Asn149 and Gly153. Models were produced using NGL viewer^54^ and the relevant Protein Data Bank entries.

#### 3.3.2. Transport Activities of NupC Cysteine Mutants

Using an assay for NupC-mediated uptake of [^14^C]uridine into energized *E. coli* whole cells,^40^ transport activities of cysteine mutants were compared to that for WT (Figure 4). Transport by WT was concentration-dependent and saturable, and the resultant curve was fitted to obtain values for the affinity of initial rate transport (*K*_m_^app^) and maximal velocity (*V*_max_) of 22.2 ± 3.7 *μ*M and 13.5 ± 0.8 nmol (mg of cells)^−1^ (15 s)^−1^, respectively. According to data obtained over a range of time points with the lowest (5 *μ*M) and highest (100 *μ*M) concentrations of [^14^C]uridine used with WT NupC and mutant S142C, 15 s is in the linear range and not approaching equilibrium (Figure S11). A time point of 15 s is one that has routinely been used for transport assays in energized whole cells with a range of similar secondary active bacterial transporters, and this has proven to be in the initial linear range. Also, *K*_m_^app^ values in the low micromolar range are expected in energized whole cells, which is consistent with values obtained for other bacterial secondary active transporter substrates, for example, [^3^H]galactose with GalP (42−59 μM),^67^ [^14^C]uridine and [^14^C]adenosine with NupG (23.6 ± 3.1 and 20.6 ± 3.4 *μ*M, respectively),^42^ [^14^C]phenyl-thio-*β*-D-glucuronide and [^14^C]-*p*-acetamidophenyl-*β*-D-glucuronide with GusB (198 ± 69 and 36 ± 4 *μ*M, respectively),^68^ [^3^H]myo-inositol with IolT (111 ± 22 *μ*M),^69^ [^14^C]allantoin with PucI (24 ± 3 *μ*M),^70^ and [^3^H]-L-glutamate with GltP (22.6 ± 5.5 μM).^71^ Mutant S142C had a *K*_m_^app^ value (18.6 ± 2.5 *μ*M) that was not significantly different from that of WT, while its *V*_max_ [10.7 ± 0.4 nmol (mg of cells)^−1^ (15 s)^−1^] was slightly lower (Figure 4). Treatment with the membrane-permeable thiol-specific reagent MTSEA reduced the rate of S142C-mediated uptake of [^14^C]uridine by around 50%.^66^ Mutants G146C and E149C both had uptake levels similar to those obtained for cells with no amplified expression of any transport protein, as demonstrated using cells transformed with plasmid pTTQ18 containing no transporter gene insert (Figure 4). This background uptake into negative control cells, which does fit to the Michaelis−Menten curve, is presumed to originate from naturally occurring levels of potentially multiple endogenous proteins capable of transporting or binding uridine. Measurements of uptake of [^14^C]adenosine and [^14^C]uridine after 15 s and 2 min by cells transformed with empty pTTQ18 compared with cells transformed with NupC-expressing plasmid pGJL16 (Figure S12) confirm the background levels of uptake seen in Figure 4. Hence, mutants G146C and E149C had both lost the ability to transport [^14^C]uridine, even up to a concentration of 400 *μ*M. Loss of transport activity was not due to a decrease in the level of protein expression, because Western blot analysis of membrane preparations from the same cells used in the transport assay demonstrated comparable levels of expression for WT NupC and the three mutants (Figure 4). Analysis of transport activity by the same NupC mutants expressed in *X. laevis* oocytes produced results broadly consistent with observations made in *E. coli* whole cells. S142C was functional and inhibited by PCMBS; G146C was non-functional, and E149C was borderline functional with a flux too low to characterize.^72^ The next consideration was whether mutations G146C and E149C had affected just transport activity or the ability of NupC to recognize and bind a substrate. The transport defective mutants may have lost the ability to undergo a conformational change required to complete the transport cycle but retained nucleoside binding activity.

**Figure 4.**
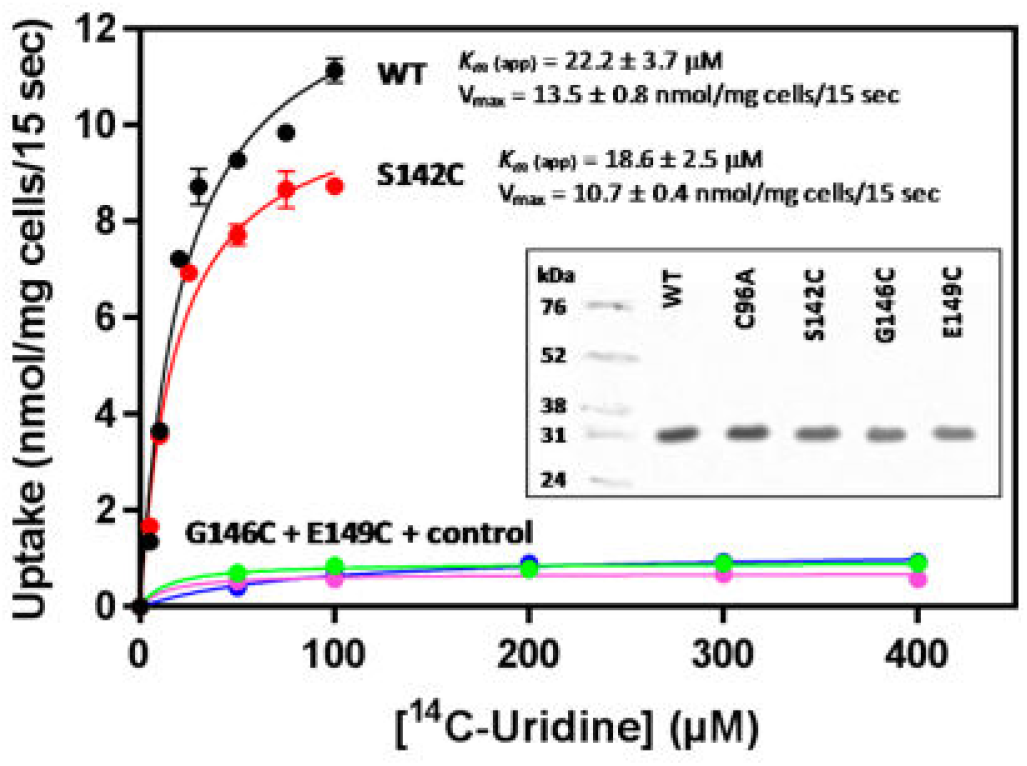
Transport activities of NupC cysteine mutants. Concentration dependence of uptake of [^14^C]uridine after 15 s by transport energized *E. coli* BL21(DE3) cells harboring plasmid pTTQ18 with no transporter gene insert (*blue*) or plasmid pGJL16 for expressing WT NupC (*black*) or constructs for expressing mutants S142C (*red*), G146C (*green*), and E149C (*pink*) following growth in minimal medium and induction with IPTG. For WT and S142C, data points represent the average of duplicate measurements. Kinetic parameters for WT and S142C were obtained by fitting data to the Michaelis− Menten equation using GraphPad Prism. The inset is a Western blot analysis of membrane preparations from the same cells used in the transport assay and also membranes from cells expressing cysteineless NupC (C96A).

#### 3.3.3. Characterization of Nucleoside Binding by NupC Cysteine Mutants Using CP-MAS Solid-State NMR

We next investigated whether the different transport activities of NupC mutants were a consequence of impaired transport function, a decrease in the affinity for the nucleoside-binding site, or both. Purified NupC was not available, and nucleosides absorb wavelengths of radiation in regions that can hamper binding measurements using techniques such as fluorescence spectroscopy and near-ultraviolet circular dichroism spectroscopy.^73,74^ Analysis of nucleoside binding was therefore conducted using ^13^C CP-MAS NMR spectroscopy measurements on the proteins with amplified expression in un-energized *E. coli* inner membranes (Figure S10). These experiments used and expanded on methods already developed to observe and quantify binding of [1′-^13^C]uridine to WT NupC.^46 13^C CP-MAS NMR spectra of NupC-enriched membranes containing [1′-^13^C]uridine exhibit a peak from the labelled substrate at a chemical shift position of ~91 ppm. At high substrate concentrations (≥5 mM), the peak intensity at a defined Hartmann−Hahn contact time reflects both specific interactions of uridine with NupC and interactions at other, undefined sites within the membranes. Fractions of total peak intensity that reflect specific and undefined interactions can be determined by displacing all of the specifically bound [1′-^13^C]uridine with a non-labelled competitor.^46^ In the work presented here, displacement NMR spectra representing only [1′-^13^C]uridine interacting specifically with NupC were obtained by the subtraction method described in Materials and Methods. Transport proteins can display relatively weak binding for their substrates to accommodate transient interactions that take place during compound recognition and the transport cycle. Apparent dissociation constants (*K*_d_^app^ values) in the low millimolar range are therefore expected for substrate binding in non-energized native membranes. This is consistent with similar solid-state NMR measurements on other bacterial secondary active transporters, including GalP, GusB, NupG, ZitB, and GltP.^42,46,71,75–79^

Normalized ^13^C CP-MAS displacement spectra were recorded for membrane preparations containing 5 mM [1′-^13^C]uridine and 80 nmol of WT NupC or mutants S142C, G146C, and E149C (Figure 5A). The background signal from natural abundance ^13^C in the membranes was removed by subtraction, and each spectrum shows only a single peak at 91 ppm representing the labelled substrate displaced from the membranes by 40 mM unlabelled uridine. Peaks representing interactions of [1′-^13^C]uridine with WT and S142C are considerably more intense than those for interactions with G146C and E149C. Peak intensities in displacement spectra are related to the specific association of [1′-^13^C]uridine and NupC only if the labelled substrate is not displaced from other, undefined sites by unlabelled uridine. When control membranes that contain NupC at the natural abundance level of <1% of total membrane protein were used, specific binding of [1′-^13^C]uridine to NupC was negligible compared with the undefined interactions. No peaks were observed in the displacement spectrum from control membranes, confirming that [1′-^13^C]uridine was not displaced from sites other than with NupC. Initial visual inspection of peak intensities in displacement spectra suggested that the affinity of uridine for G146C and E149C was somewhat different from that for WT. It is not possible to obtain quantitative information from single spectra, so further experiments were necessary to establish how the mutations had affected binding affinity.

**Figure 5.**
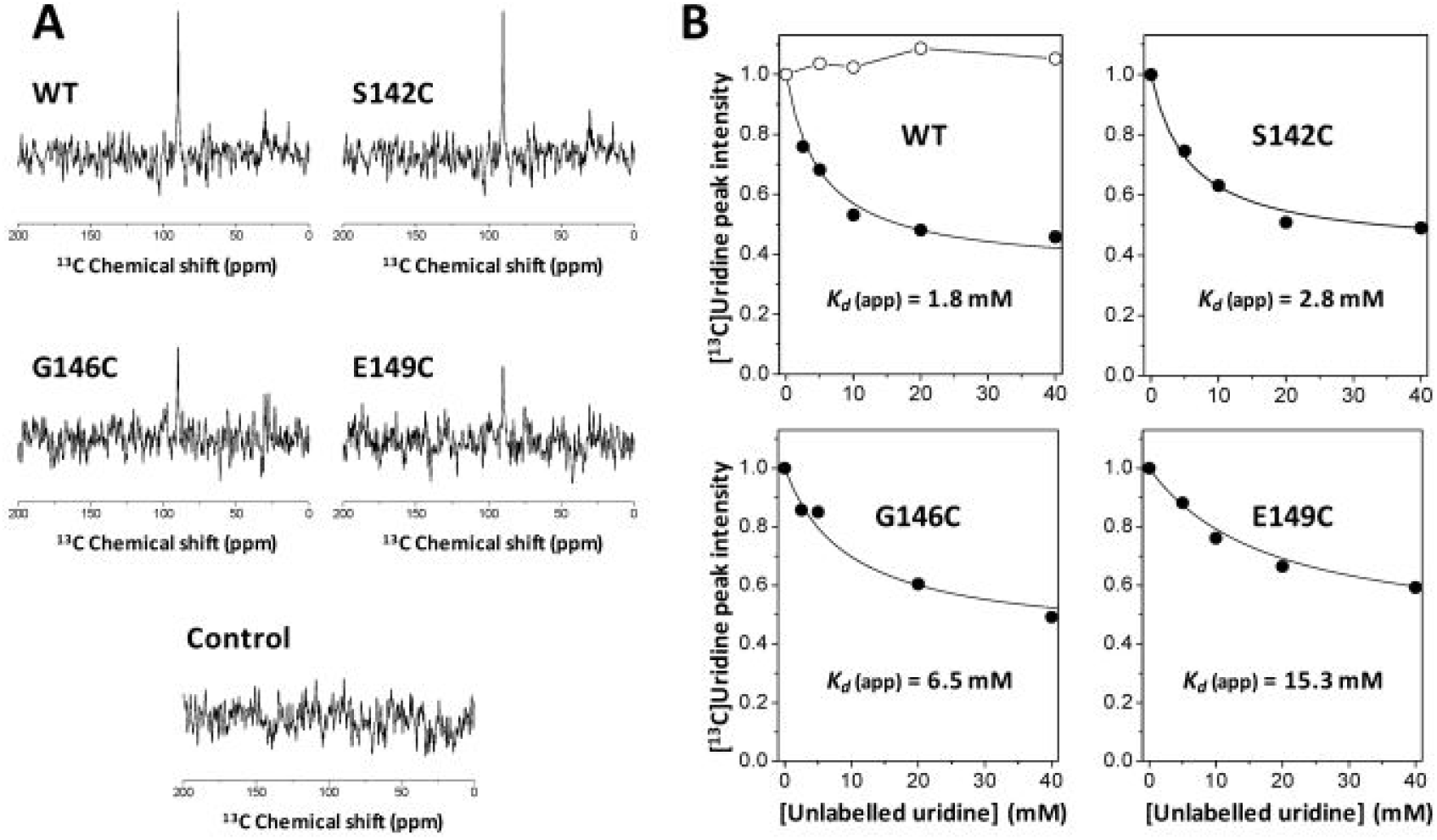
Relative affinities of uridine binding to wild-type NupC and mutants in native inner membranes. (A) Displacement ^13^C CP-MAS NMR spectra obtained from un-energized *E. coli* inner membranes containing 5 mM [1′-^13^C]uridine and 80 nmol of WT NupC or mutants S142C, G146C, and E149C. Displacement spectra were obtained by the procedure described in Materials and Methods, in which spectra of each membrane sample were obtained before and after the addition of 40 or 80 mM unlabelled uridine to displace the labelled substrate. Each spectrum was obtained using a Hartmann−Hahn contact time of 10 ms and was the result of the accumulation of 6144 scans. Spectra from membranes with amplified expression of WT, S142C, G146C, or E149C all exhibit a peak at 91 ppm from the substrate interacting specifically with NupC. Also shown is a control displacement spectrum from membranes containing natural abundance levels of NupC, which indicates that the peak intensity was not displaced. Difference spectra were normalized to the intensity of the lipid peak at 32 ppm in the original spectra. (B) Curves of ^13^C CP-MAS peak intensities for [1′-^13^C]uridine in membranes containing WT NupC or mutants S142C, G146C, and E149C at increasing concentrations of unlabelled D-uridine. Peak intensities (●) were obtained from spectra recorded using a Hartmann−Hahn contact time (*t*_c_) of 2 ms after 4096 scans. Also shown in the plot for WT NupC are peak intensities for [1′-^13^C]uridine in the presence of increasing concentrations of unlabelled L-uridine (○). Solid lines are curves of best fit calculated according to Equation 2 corresponding to the values of the apparent *K*_d_ shown in each plot.

Dissociation constants for substrates of bacterial secondary transporters have been estimated previously using ^13^C CP-MAS NMR measurements of displacement by a competitive substrate.^42,46,71,76,78^ When the rate of dissociation of the substrate is slow, the NMR peak intensity for a labelled substrate is related to the concentration of the competitive substrate by an inverse rectangular hyperbolic function. The total peak intensity (*I*_T_) from a labelled ligand (L*) is a function of contact time (*t*_c_) and the concentration of unlabelled competitor ligand (L). In the case of a labelled ligand such as [1′-^13^C]uridine, *I*_T_ has components *I*_S_ from the specifically bound substrate and *I*_NS_ from the remaining nonspecific or undefined interactions of the substrate. For example

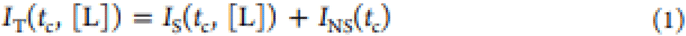

If total peak intensity is normalized to unity when the concentration of unlabelled ligand [L] is zero, i.e., *I*_T_(*t*_c_,[L]_0_) = 1, it follows that

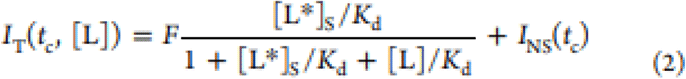

where

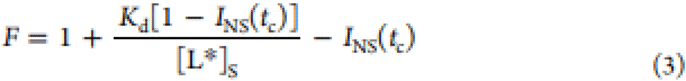

*I*_NS_(*t*_c_) is measured as the peak intensity remaining after the specifically bound ligand has been fully displaced by the unlabelled ligand. Values of *K*_d_ and [L*]_S_ can be estimated by nonlinear least-squares fitting of Equation 2 to NMR peak intensity curves measured in a titration experiment. Equations 1–3 assume that dilution effects because of titrating unlabelled substrate solution into the membranes are negligible. Equations 1–3 hold only when a linear relationship exists between *I*_S_ and the concentration of the specifically bound ligand, which occurs only when the rate of ligand dissociation is slow.^46^ When the off rate of the ligand is not known, the measured *K*_d_ represents a lower limit value and further experiments are necessary. It is also assumed that component *I*_NS_(*t*_c_) from the non-specifically bound ligand remains constant and is not diminished by addition of the unlabelled substrate. The validity of this assumption was supported by the lack of signal observed in the displacement spectrum from the control membranes (Figure 5A). Dissociation constants for interaction between [1′-^13^C]uridine and NupC were estimated from curves of ^13^C NMR peak intensities at 91 ppm measured after titrating unlabelled uridine into the membrane sample to a final concentration of 40 mM (Figure 5B). A decrease in peak intensity with increasing concentrations of unlabelled substrate was observed for WT NupC, for which curve fitting (Figure 5B) yielded an apparent *K*_d_ of 1.8 mM and a concentration for specifically bound [1′-^13^C]uridine of 3.5 mM (i.e., 70% of the total substrate concentration). The *K*_d_ value is slightly lower than the value of 2.6 mM measured previously.^46^ Similar displacement curves for the mutants produced apparent *K*_d_ values of 2.8, 6.5, and 15.3 mM for S142C, G146C, and E149C, respectively (Figure 5B). In all three cases, best fitting curves corresponded to a concentration of specifically bound uridine of 3.5 mM. In a control experiment with WT NupC containing [1′-^13^C]uridine and the unlabelled non-substrate L-uridine (Figure 5B), the peak intensity did not decrease upon titration with the stereoisomer. This experiment confirmed that the decrease in peak intensity produced by addition of the D-uridine substrate was a direct result of displacing [1′-^13^C]uridine from NupC.

*K*_d_ values obtained in the titration experiments serve only as estimates because it is not known whether the dissociation rates of uridine are slow enough for Equation 2 to hold. Binding affinities and dissociation rate constants (*k*_off_) for [1′-^13^C]uridine in NupC membranes were therefore estimated using variable contact time ^13^C CP-MAS NMR as an alternative method. Peak intensity curves were measured for the labelled substrate in the membranes over a range of contact times of ≤10 ms. These were compared with curves calculated for different *K*_d_ and *k*_off_ values to determine the combination of values that minimized the χ^2^ function. Experimental peak intensity curves for membranes containing 5 mM [1′-^13^C]uridine and 80 nmol of WT NupC are shown in Figure 6. The closest agreement between experimental and simulated curves was for a *K*_d_ of 2.6 mM and a *k*_off_ of 333 s^−1^. It should be noted, however, that curves for *K*_d_ values between 0.5 and 5.5 mM fell within the experimental error. Similar experiments with the mutants yielded χ^2^ minima for *K*_d_ values of 3.1, 11.9, and 13.4 mM for S142C, G146C, and E149C, respectively (Figure 6). In each case, *k*_off_ decreased at values of 300−400 s^−1^. There is reasonable agreement between the *K*_d_ values obtained by the variable contact time and displacement methods, although values for G146C (6.5 and 11.9 mM) are somewhat different. The quality of NMR data that can be obtained in a practicable experimental time (12 h) undoubtedly limits the precision and accuracy of measurements, so caution must be exercised in drawing conclusions from the absolute values. Both NMR methods of analysis identify the same trend in binding affinity of [1′-^13^C]uridine for WT NupC and the mutants (i.e., WT > S142C > G146C > E149C), however, and conclusions about relative substrate affinities can be made with greater confidence. In summary, mutation S142C appears to have only a weak effect on the affinity for uridine, while G146C and E149C decrease the affinity for uridine. The latter two mutants are still able to bind uridine, however, while the transport of [^14^C]uridine in whole *E. coli* cells was abolished (Figure 4). Gly146 and Glu149 therefore appear to be important for uridine binding but are essential for uridine transport.

**Figure 6.**
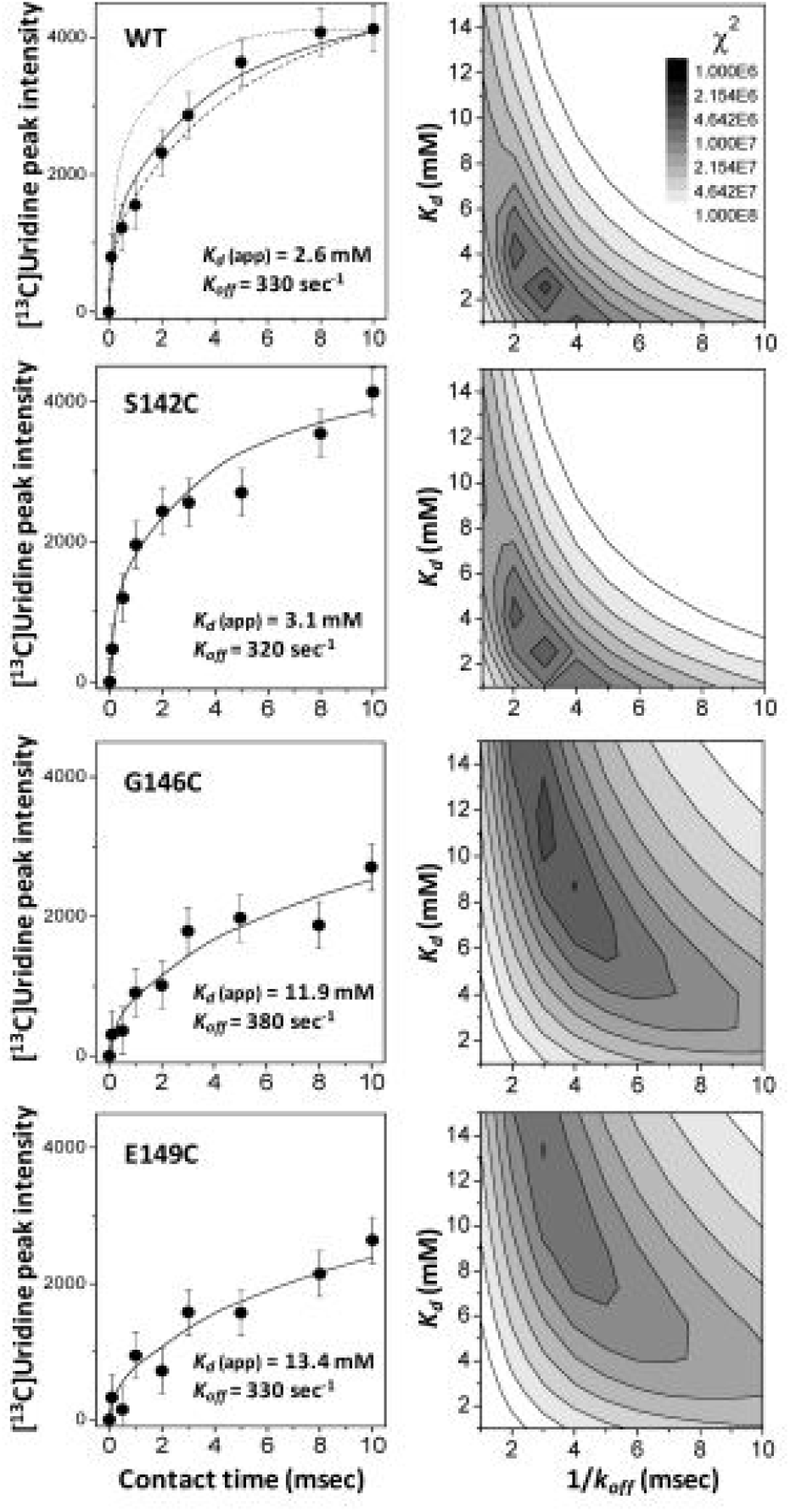
Variable contact time analysis of uridine binding to wild-type NupC and mutants in native inner membranes. Experimental curves and simulations of ^13^C peak intensities at 91 ppm from variable contact time displacement spectra of 5 mM [1′-^13^C]uridine in *E. coli* inner membranes containing 80 nmol of WT NupC or mutants S142C, G146C, and E149C. Displacement spectra were obtained as described in Materials and Methods using 40 mM unlabelled uridine as the displacing substrate. The points of the curve (left, circles) represent specific binding and were measured as the decrease in peak intensity at each contact time after the labelled substrate was displaced by unlabelled uridine. Error bars represent the level of noise in the spectra. Simulated curves were computed for different combinations of *K*_d_ and dissociation rate constant *k*_off_ using the Monte Carlo method described previously.^46^ The goodness of fit of each curve to the experimental data was assessed from the function χ^2^ = ∑_*i*_[(*S_i_* − *E*)/*σ_ii_*]^2^. The contour plot (right) is shaded according to the χ^2^ values (shown in the key) obtained for different combinations of *K*_d_ and *k*_off_. For example, in the case of WT, the χ^2^ minimum (denoted by the darkest region on the contour plot) was obtained for a *K*_d_ of 2.6 mM and a *k*_off_ of 3.1 ms. The curve corresponding to the χ^2^ minimum is shown on the intensity plot (right) as a solid line, alongside curves for *K*_d_ values of 0.5 mM (dotted line) and 5.5 mM (dashed line) for comparison. Fixed simulation parameters were as follows: ^1^H *T*_1ρ_ = 100 ms for free substrate, ^1^H *T*_1ρ_ = 8 ms for bound substrate, *R*_HC_ = 5 × 10^−3^ s^−1^, protein content = 80 nmol, and substrate concentration = 5 mM.

Experiments were also conducted to compare the ability of WT NupC and the mutants to bind the uridine-competitive substrate thymidine. Interaction of thymidine with NupC was observed indirectly through displacement of 5 mM [1′-^13^C]-uridine with 50 mM unlabelled thymidine (Figure 7). The peak intensity for WT was reduced by ~50%, similar to that observed when specifically bound [1′-^13^C]uridine was fully displaced by 40 mM unlabelled uridine (Figure 5B). From these spectra, it is reasonable to conclude that all, or most, of the labelled uridine is displaced from WT NupC by 50 mM thymidine, as would be expected for competitive substrates having a similar affinity for NupC. Spectra for S142C and G146C also show an ~50% drop in peak intensity for [1′-^13^C]uridine upon addition of thymidine (Figure 7), again suggesting that uridine and thymidine have similar affinities for these mutants. By contrast, the spectrum for E149C shows little or no loss of peak intensity following addition of thymidine (Figure 7). Of the three mutants, E149C has the lowest affinity for uridine (*K*_d_ > 15 mM) and an inability of 50 mM thymidine to displace specifically bound uridine, indicating that this mutant has an even lower affinity for thymidine.

**Figure 7.**
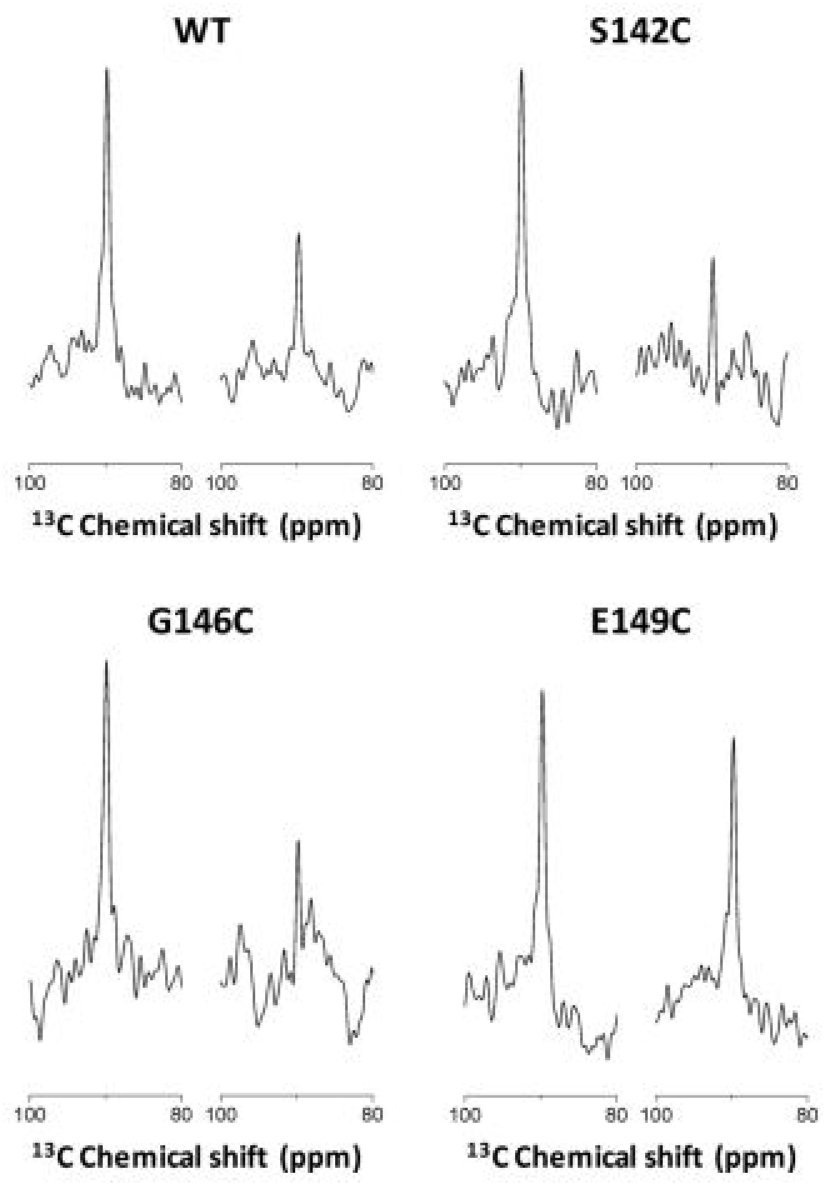
Competitive binding of thymidine to wild-type NupC and mutants in native inner membranes. Regions of ^13^C CP-MAS NMR spectra of [1′-^13^C]uridine in *E. coli* inner membranes containing WT NupC or mutants S142C, G146C, and E149C. Spectra show the peak at 91 ppm from the labelled substrate in the absence (*left*) or presence (*right*) of 50 mM unlabeled thymidine. Spectra were recorded using a Hartmann−Hahn contact time (*t*_c_) of 2 ms after 6144 scans.

A further experiment investigated the effect of cysteine reactive iodoacetate on [1′-^13^C]uridine binding by mutant E149C and thus the proximity of Glu149 to the uridine-binding site (Figure 8). Control ^13^C CP-MAS measurements were first performed on membranes with amplified expression of cysteineless NupC (C96A), whereby [1′-^13^C]uridine binding was displaced by 50 mM unlabelled uridine but was not affected by pre-treatment with 10 mM iodoacetate. In the case of membranes with amplified expression of E149C, [1′-^13^C]-uridine binding was displaced by 50 mM unlabelled uridine and also blocked by pre-treatment with 10 mM iodoacetate. This result confirms the direct involvement of Glu149 in uridine binding by NupC.

**Figure 8.**
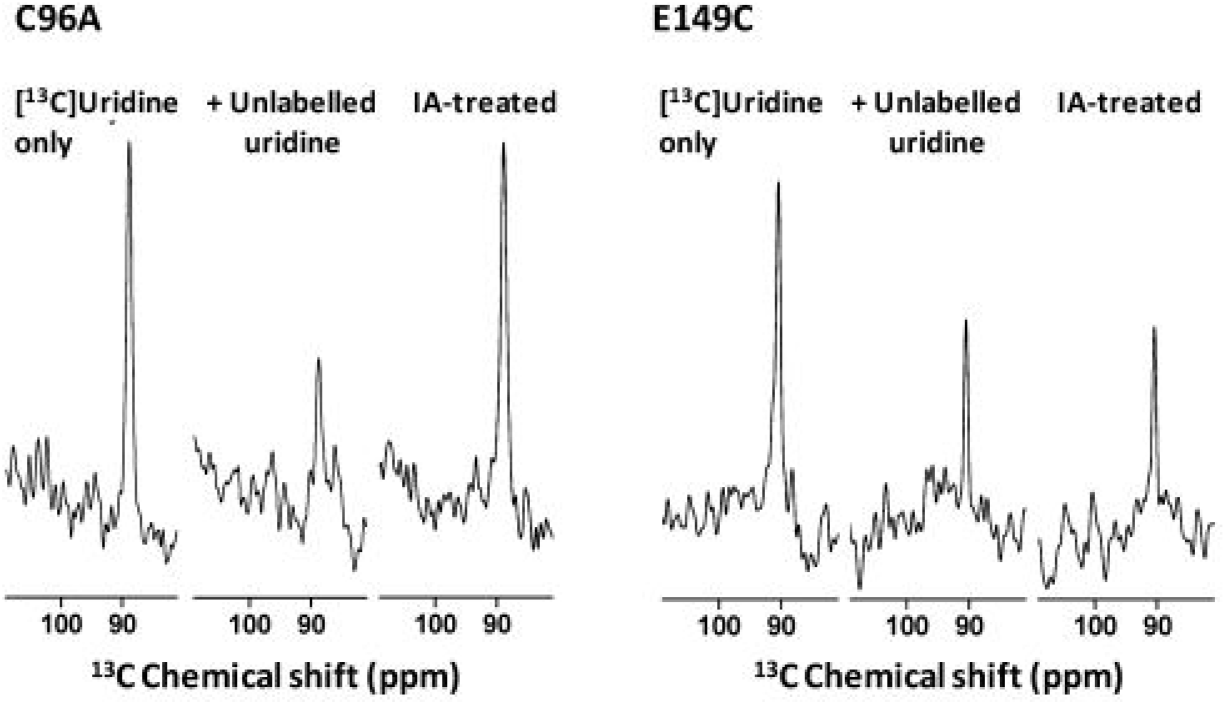
Elimination of binding of uridine to NupC mutant E149C in native inner membranes by treatment with iodoacetate. Expansions of the signals from ^13^C CP-MAS NMR spectra are shown for binding of 5 mM [1′-^13^C]uridine in inner membranes with amplified expression of NupC mutants C96A (cysteineless NupC) and E149C for samples in the absence of any other additions and after pre-treatment with 10 mM iodoacetate (IA) and/or competition with 50 mM unlabelled uridine. For pre-treatment with iodoacetate, membranes containing NupC protein at a concentration of ~0.1 mM were incubated with the following successive additions at 4 °C: 2 mM DTT for 30 min, 10 mM iodoacetate for 1 h in the dark, 20 mM DTT for 15 min (to mop up excess iodoacetate), and then 5 mM [1′-^13^C]uridine for 30 min. Spectra were recorded using a Hartmann−Hahn contact time (*t*_c_) of 2 ms after 7200 and 6000 scans for C96A and E149C, respectively.

### 3.4. Explanation of Results with Respect to an Elevator-Type Mechanism of Alternating Access Membrane Transport in NupC

A homology model of a NupC protomer based on the crystal structure of vcCNT in complex with uridine predicts that mutated residues Ser142, Gly146, and Glu149 are located at the tip of HP1 and close to the putative nucleoside-binding site (Figure 9A). While the position corresponding to Ser142 is conserved as asparagine in vcCNT, nwCNT, and hCNTs and involved in coordinating the sodium ion in these proteins, the hydroxyl on Ser142 is not essential for substrate binding or transport activity in NupC. The NMR results suggest that mutations G146C and E149C have disrupted interactions between the substrate and its binding site. The abolished transport activity by G146C and E149C also suggests that these mutations have blocked substrate binding interactions and/or conformational changes required for completing the transport cycle.

**Figure 9.**
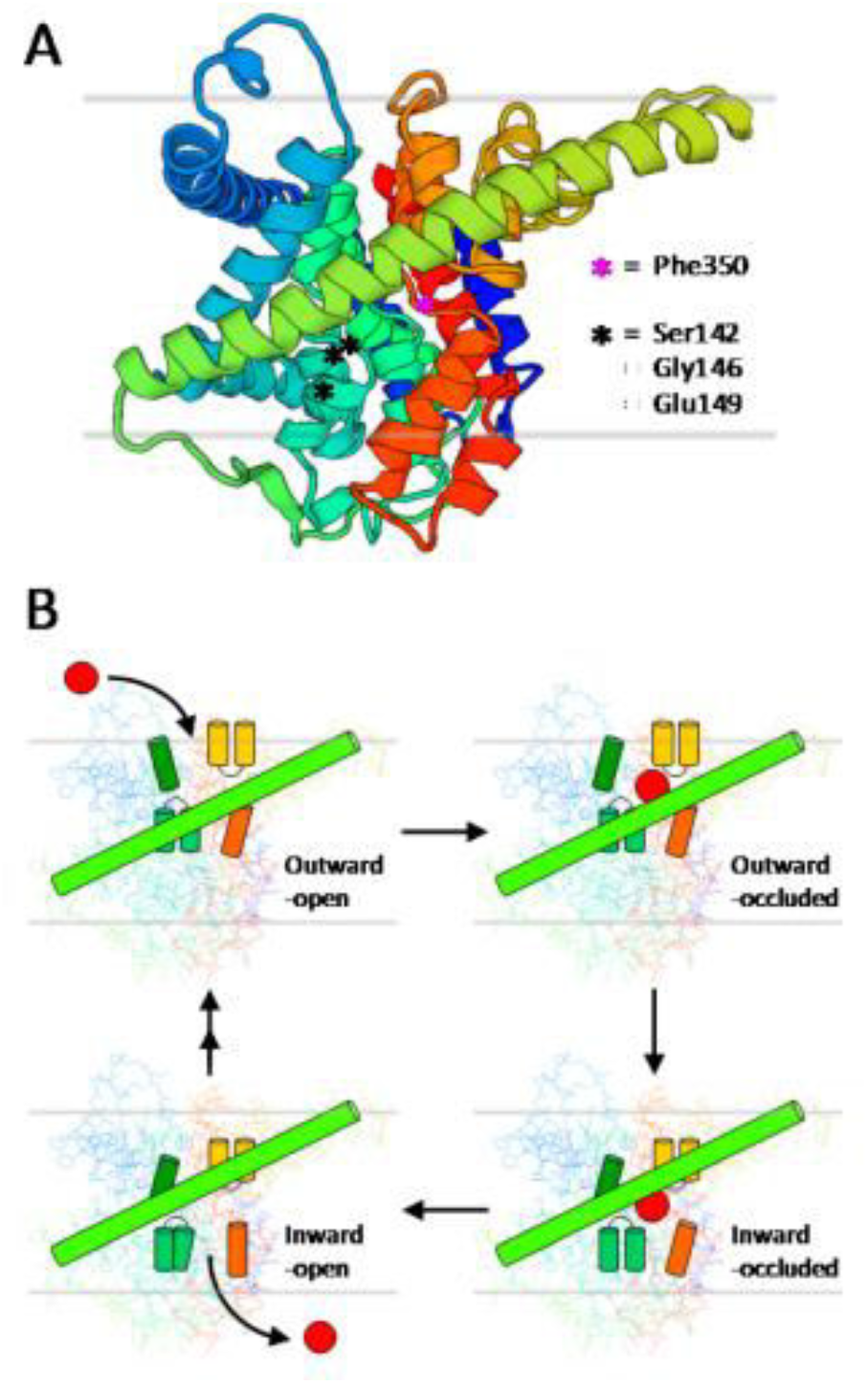
Homology model of a NupC protomer and cartoon representation of a hypothetical elevator-type mechanism of concentrative nucleoside transport by NupC. (A) Homology model of a NupC protomer based on the crystal structure of vcCNT in complex with uridine (Protein Data Bank entry 4PD6) produced using SWISS-MODEL (https://swissmodel.expasy.org/).^55^ The model includes a blue N-terminus and a red C-terminus, and the positions of mutated HP1 residues Ser142, Gly146, and Glu149 are indicated. The position of Phe350 that corresponds to the crucial vcCNT TM7b residue Phe366 is also indicated. (B) Cartoon representation of a hypothetical elevator-type mechanism of alternating access concentrative nucleoside transport by a NupC protomer. The external nucleoside (*red circle*) binds to an outward-open state that undergoes conformational change to an outward-occluded state in which the nucleoside packs between the tips of HP1 (*mid green*) and HP2 (*light orange*), TM4b (*dark green*), and TM7b (*dark orange*) behind the center of TM6 (*light green*). Elevator-like movement of the nucleoside-binding site (including HP1, HP2, TM4b, and TM7b) across TM6 produces an inward-occluded state that undergoes conformational change to an inward-open state from which the nucleoside is released to inside the cell. The outward-open state is then recovered. TM4b and HP1b act as gates for the outward-facing and inward-facing end states, respectively. The horizontal grey lines represent the approximate outer and inner limits of the lipid bilayer.

On the basis of sequence homology, predictions of transmembrane helices, homology modeling, and the results of substrate binding and transport experiments, we have no reason to believe that NupC does not also adopt an elevator-type mechanism of alternating access membrane transport similar to that described for vcCNT and nwCNT.^24,28,29^ In a hypothetical transport cycle for NupC (Figure 9B), the external nucleoside binds to an outward-open state that undergoes a conformational change to an outward-occluded state in which the nucleoside packs between the tips of HP1 and HP2, TM4b, and TM7b behind the center of TM6. Elevator-like movement of the nucleoside-binding site (including HP1, HP2, TM4b, and TM7b) across TM6 produces an inward-occluded state that undergoes a conformational change to an inward-open state from which the nucleoside is released. The outward-open state is then recovered, possibly via multiple intermediate conformations. TM4b and HP1b act as gates for the outward-facing and inward-facing end states, respectively. It is therefore easy to envisage how mutations G146C and E149C at the tip of HP1 could block conformational changes in the transport cycle.

## 4. Conclusion

*E. coli* CNT family protein NupC shares the highest degree of sequence homology with crystallographically defined vcCNT and nwCNT (30.3 and 32.3% identical and 28.8 and 27.3% highly similar, respectively; combined 59.1 and 59.6%, respectively) and then with the human proteins in the following order: hCNT3 > hCNT1 > hCNT2. Regions of NupC corresponding to vcCNT structural domains involved in nucleoside binding (HP1, TM4, HP2, and TM7) contain highly conserved motifs. Of nine vcCNT residues involved in direct binding interactions with nucleosides, seven are identically conserved in NupC and the other two are highly similar. The corresponding residues in NupC are Gly146, Gln147, Ser148, Glu149, Ile181, Glu321, Phe350, Asn352, and Ser355. A large majority of these residues are also conserved in hCNTs. The four sodium-binding site residues in vcCNT are not conserved in NupC, reflecting their differences in cation selectivity. Based on mutations that change hCNT1 from a pyrimidine-specific to a purine-specific transporter, three mutations (Q147M, S180T, and I181V) are predicted to have the same effect on NupC.

Analysis of the NupC sequence by membrane topology prediction tools suggested nine putative transmembrane helices (or eight with a partial helix). The N-terminus was predicted to be periplasmic or cytoplasmic, but the distribution of positively charged residues was more consistent with a periplasmic N-terminus as it is in vcCNT. The putative locations of transmembrane helices in NupC based on sequence homology and the structure of vcCNT all showed some overlap with those coming from prediction tools, but there were significant differences in their lengths and in their start and end positions. The extra “ninth” helix in NupC given by prediction tools appears in a region corresponding to IH3 in vcCNT. In comparison, analysis of the vcCNT sequence by membrane topology prediction tools suggested ten or eight transmembrane helices with periplasmic N- and C-termini. While eight of these transmembrane helices showed some overlap with those in the structure of vcCNT, significant differences in their lengths and in their start and end positions emphasized the challenges in predicting transmembrane helices in CNTs.

Cysteine scanning mutagenesis of NupC investigated roles of specific residues in nucleoside binding and/or transport activity. WT NupC had an apparent transport affinity (*K*_m_^app^) for[^14^C]uridine of 22.2 ± 3.7 μM in energized *E. coli* whole cells and an apparent binding affinity (*K*_d_^app^) for [1′-^13^C]uridine of 1.8−2.6 mM in non-energized inner membranes. The position corresponding to Ser142 is occupied by asparagine in vcCNT, nwCNT, and hCNTs; in the structure of vcCNT, this is involved in coordinating the sodium ion. Consistent with this, NupC mutant S142C retained transport and binding affinities similar to those of WT. Gly146 is conserved in vcCNT, nwCNT, hCNT2, and hCNT3 and corresponds with Ser319 in hCNT1 that when mutated to glycine allows purine nucleoside transport. Glu149 corresponds to Glu156 in vcCNT, which is involved in nucleoside binding, and this position is conserved as glutamate in nwCNT and hCNTs. Consistent with these observations, mutants G146C and E149C both had abolished transport activity. These mutants did retain varying degrees of partial uridine binding activity, however, with affinities decreasing in the following order: WT > S142C > G146C > E149C. Gly146 and Glu149 therefore appear to be important for uridine binding but are essential for uridine transport. WT NupC and mutants S142C and G146C had similar respective affinities for binding uridine and thymidine. Mutant E149C had an even lower affinity for thymidine than for uridine, and treatment of E149C with iodoacetate blocked uridine binding, thus confirming the direct involvement of Glu149 in binding nucleosides.

In addition to vcCNT and nwCNT, NupC is a viable model for concentrative nucleoside transport, including the human proteins. Hence, there is a compelling requirement for high-resolution crystal structures of NupC determined in multiple conformations and in complex with different ligands. These would confirm the putative structural organization of NupC and determine if it adopts the same elevator-type mechanism of alternating access membrane transport proposed for vcCNT and nwCNT. They would also demonstrate structural differences in substrate binding and the transport cycle for proton-driven versus sodium-driven nucleoside transport. Such structures could help explain how hCNT3 can use a proton gradient as well as a sodium gradient to drive uridine transport. NupC mutants with abolished transport activity, such as G146C and E149C from this study, may be locked in a single conformation of the transport cycle and therefore potentially more tractable to crystallization than the WT protein. Crystal structures of NupC could be combined with sub-angstrom resolution structural information about nucleoside conformation and interactions in the substrate-binding site using isotope labels and solid-state NMR methods that measure internuclear distances and torsion angles.^45,58,77,80–82^ Also, construction of further site-specific mutants and their characterization using a range of computational, chemical, biochemical, and biophysical methods should be pursued.

## Acknowledgements

The authors thank David Middleton (Lancaster University, Lancaster, UK) for technical assistance with solid-state NMR experiments at 400 MHz.

