## Supporting Information for "Elevator Mechanism of Alternating Access in the *Escherichia coli* Concentrative Nucleoside Transporter NupC"

(Tables S1 and S2, Figures S1-S12)

### Elevator Mechanism of Alternating Access in the *Escherichia coli* Concentrative Nucleoside Transporter NupC


Lijie Sun and Simon G. Patching*

School of Biomedical Sciences and Astbury Centre, University of Leeds, Leeds, UK

**Table S1. Sequence homology in concentrative nucleoside transporters.** Amino acid sequences of NupC from *Escherichia coli* (P0AFF2), vcCNT from *Vibrio cholera* (Q9KPL5), nwCNT from *Neisseria wadsworthii* (G4CRQ5) and human concentrative nucleoside transporters hCNT1 (O00337), hCNT2 (O43868) and hCNT3 (Q9HAS3) were taken from the UniProt KnowledgeBase (http://www.uniprot.org/) and separately aligned with each other using the online multiple sequence alignment tool Clustal Omega (http://www.ebi.ac.uk/Tools/msa/clustalo/).[1] The percentage sequence homology is given in terms of identical residues (*red*), highly similar residues (*blue*) and the combined total of these (*black*).

| | NupC | | | vcCNT | | | nwCNT | | | hCNT1 | | | hCNT2 | | | hCNT3 | | |
|---|---|---|---|---|---|---|---|---|---|---|---|---|---|---|---|---|---|---|
| **NupC** | | | | 30.3 | 28.8 | 58.1 | 32.3 | 27.3 | 59.6 | 27.0 | 26.3 | 53.3 | 22.3 | 31.3 | 53.5 | 25.3 | 29.3 | 54.5 |
| **vcCNT** | 30.3 | 28.8 | 59.1 | | | | 66.5 | 18.2 | 84.7 | 36.4 | 26.8 | 63.2 | 36.1 | 26.6 | 62.7 | 38.8 | 25.4 | 64.2 |
| **nwCNT** | 32.3 | 27.3 | 59.6 | 66.5 | 18.2 | 84.7 | | | | 35.5 | 25.6 | 61.1 | 34.6 | 25.9 | 60.5 | 34.6 | 27.3 | 61.9 |
| **hCNT1** | 27.0 | 26.3 | 53.3 | 36.4 | 26.8 | 63.2 | 35.5 | 25.6 | 61.1 | | | | 63.3 | 17.7 | 81 | 43.9 | 23.4 | 67.3 |
| **hCNT2** | 22.3 | 31.3 | 53.5 | 36.1 | 26.6 | 62.7 | 34.6 | 25.9 | 60.5 | 63.3 | 17.7 | 81 | | | | 42.1 | 24.3 | 66.4 |
| **hCNT3** | 25.3 | 29.3 | 54.5 | 38.8 | 25.4 | 64.2 | 34.6 | 27.3 | 61.9 | 43.9 | 23.4 | 67.3 | 42.1 | 24.3 | 66.4 | | | |

**Table S2. Amino acid compositions of concentrative nucleoside transporters.** Amino acid sequences of NupC from *Escherichia coli* (P0AFF2), vcCNT from *Vibrio cholera* (Q9KPL5), nwCNT from *Neisseria wadsworthii* (G4CRQ5) and human concentrative nucleoside transporters hCNT1 (O00337), hCNT2 (O43868) and hCNT3 (Q9HAS3) were taken from the UniProt KnowledgeBase (http://www.uniprot.org/) and analysed by the online ExPASy tool ProtParam (http://web.expasy.org/protparam/).[2] The contents of individual amino acids and of groupings of amino acids with similar physicochemical properties in each nucleoside transporter were compared with overall average values in 235 secondary transport proteins from *Escherichia coli* and in 336 human secondary transport proteins.[3,4] The groupings of amino acids with similar physicochemical properties are: hydrophobic (alanine, isoleucine, leucine, phenylalanine and valine); positively charged at physiological pH (arginine and lysine); negatively charged at physiological pH (aspartic acid and glutamic acid); hydroxyl-containing (serine and threonine); amido-containing (asparagine and glutamine).

| Amino acid | *E. coli* | Human | NupC | vcCNT | nwCNT | hCNT1 | hCNT2 | hCNT3 |
|---|---|---|---|---|---|---|---|---|
| **Ala** | 10.8% | 8.2% | 9.8% | 13.2% | 13.6% | 10.3% | 9.4% | 7.7% |
| **Arg** | 3.5% | 4.2% | 2.8% | 2.6% | 2.6% | 5.4% | 3.6% | 3.9% |
| **Asn** | 2.8% | 3.0% | 4.2% | 3.3% | 2.8% | 2.3% | 3.0% | 4.3% |
| **Asp** | 2.3% | 3.1% | 1.8% | 2.1% | 2.1% | 2.8% | 2.1% | 2.6% |
| **Cys** | 1.1% | 2.1% | 0.2% | 0.9% | 0.2% | 3.1% | 3.0% | 2.0% |
| **Gln** | 2.6% | 3.4% | 2.5% | 2.1% | 2.6% | 3.7% | 3.2% | 3.0% |
| **Glu** | 2.6% | 4.4% | 4.2% | 3.8% | 4.0% | 5.2% | 5.0% | 4.9% |
| **Gly** | 8.9% | 8.0% | 7.8% | 13.2% | 11.8% | 6.6% | 7.9% | 7.4% |
| **His** | 1.4% | 1.8% | 0.8% | 0.0% | 0.5% | 1.2% | 1.1% | 2.3% |
| **Ile** | 8.1% | 6.2% | 9.8% | 6.8% | 7.5% | 5.4% | 6.1% | 6.9% |
| **Leu** | 13.6% | 12.6% | 12.8% | 13.0% | 12.5% | 14.3% | 12.0% | 10.6% |
| **Lys** | 2.9% | 3.8% | 3.8% | 3.3% | 3.5% | 2.9% | 4.3% | 3.8% |
| **Met** | 4.0% | 2.8% | 4.5% | 4.2% | 3.8% | 2.5% | 2.6% | 3.3% |
| **Phe** | 6.1% | 5.8% | 7.8% | 7.3% | 5.2% | 6.5% | 6.8% | 6.8% |
| **Pro** | 4.0% | 5.1% | 1.8% | 4.0% | 3.1% | 3.5% | 3.2% | 3.6% |
| **Ser** | 6.6% | 7.5% | 9.2% | 6.4% | 7.1% | 8.0% | 7.8% | 8.8% |
| **Thr** | 5.5% | 5.5% | 2.8% | 3.3% | 3.3% | 4.2% | 6.2% | 5.5% |
| **Trp** | 2.1% | 1.6% | 0.8% | 1.2% | 0.9% | 2.2% | 2.0% | 2.3% |
| **Tyr** | 2.7% | 3.2% | 2.8% | 1.9% | 2.1% | 2.0% | 2.4% | 3.0% |
| **Val** | 8.4% | 7.8% | 10.2% | 7.3% | 11.1% | 7.9% | 8.2% | 7.1% |
| **Hydrophobic** | 47.0% | 40.5% | 50.4% | 47.6% | 49.9% | 44.4% | 42.5% | 39.1% |
| **Positive** | 6.4% | 8.0% | 6.6% | 5.9% | 6.1% | 8.3% | 7.9% | 7.7% |
| **Negative** | 4.8% | 7.4% | 6.0% | 5.9% | 6.1% | 8.0% | 7.1% | 7.5% |
| **Hydroxyl** | 12.1% | 13.0% | 12.0% | 9.7% | 10.4% | 12.2% | 14.0% | 14.3% |
| **Amido** | 5.5% | 6.4% | 6.7% | 5.4% | 5.4% | 6.0% | 6.2% | 7.3% |

**Figure S1. Evolutionary relationships of NupC and concentrative nucleoside transporters.** Amino acid sequences of NupC from *Escherichia coli* (P0AFF2), vcCNT from *Vibrio cholera* (Q9KPL5), nwCNT from *Neisseria wadsworthii* (G4CRQ5) and human concentrative nucleoside transporters hCNT1 (O00337), hCNT2 (O43868) and hCNT3 (Q9HAS3) were taken from the UniProt KnowledgeBase (http://www.uniprot.org/) and aligned using the online multiple sequence alignment tool Clustal Omega (http://www.ebi.ac.uk/Tools/msa/clustalo/).[1] The resultant neighbour-joining phylogenetic tree was exported in Newick format and drawn using the online tool iTOL: Interactive Tree of Life (http://itol.embl.de/).[5] Sequence homologies between NupC and the other proteins are given in terms of identical residues (*red*), highly similar residues (*blue*) and the combined total of these (*black*). These values were calculated from individual sequence alignments between NupC and the other proteins.

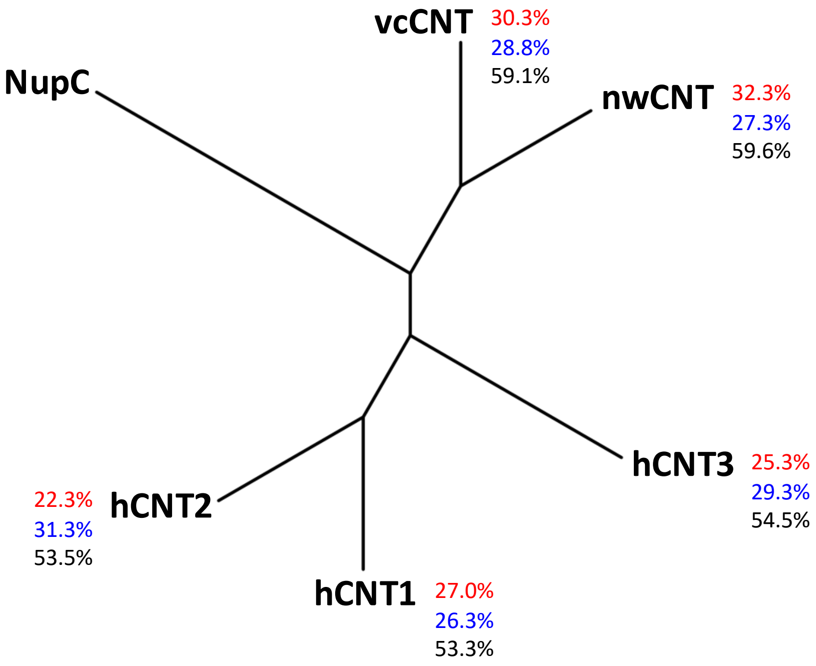

**Figure S2. Protein sequence alignment between human concentrative nucleoside transporters.** Amino acid sequences of the human concentrative nucleoside transporters hCNT1 (O00337), hCNT2 (O43868) and hCNT3 (Q9HAS3) were taken from the UniProt KnowledgeBase (http://www.uniprot.org/) and aligned using the online multiple sequence alignment tool Clustal Omega (http://www.ebi.ac.uk/Tools/msa/clustalo/).[1] Residues are coloured to indicate those that are identical (*red*) and highly similar (*blue*).

```
hCNT1   -------------------------MENDPSRRRESISLTPVAKGLENMGADFLESLEEG
hCNT2   ---------------------------MEKASGRQSIALSTVETGTVNPGLELMEKEVEP
hCNT3   MELRSTAAPRAEGYSNVGFQNEENFLENENTSGNNSIRSRAVQSREHTNTKQDEEQVTVE

hCNT1   Q-LPRSDLSPAEIRSSWSEAAPKPFSRWRNLQPALRARSFCREHMQLFRWIGTGLLCTGL
hCNT2   EGSKRTDAQGHSLGDG---LGPSTY-QRRSRWPFSKARSFCKTHASLFKKILLGLLCLAY
hCNT3   QDSPRNREH---MEDDDEEMQQKG----CLERRYDTVCGFCRKHKTTLRHIIWGILLAGY

hCNT1   SAFLLVACLLDFQRALALFVLTCVVLTFLGHRLLKRLLGPKLRRFLKPQGH--PRLLLWF
hCNT2   AAYLLAACILNFQRALALFVITCLVIFVLVHSFLKKLLGKKLTRCLKPFEN--SRLRLWT
hCNT3   LVMVISACVLNFHRALPLFVITVAAIFFVVWDHLMAKYEHRIDEMLSPGRRLLNSHWFWL

hCNT1   KRGLALAAFLGLVLWLSLDTSQR-PEQLVSFAGICVFVALLFACSKHHCAVSWRAVSWGL
hCNT2   KWVFAGVSLVGLILWLALDTAQR-PEQLIPFAGICMFILILFACSKHHSAVSWRTVFSGL
hCNT3   KWVIWSSLVLAVIFWLAFDTAKLGQQQLVSFGGLIMYIVLLFLFSKYPTRVYWRPVLWGI

hCNT1   GLQFVLGLLVIRTEPGFIAFEWLGEQIRIFLSYTKAGSSFVFGEALVKDVFAFQVLPIIV
hCNT2   GLQFVFGILVIRTDLGYTVFQWLGEQVQIFLNYTVAGSSFVFGDTLVKDVFAFQALPIII
hCNT3   GLQFLLGLLILRTDPGFIAFDWLGRQVQTFLEYTDAGASFVFGEKYKDHFFAFKVLPIVV

hCNT1   FFSCVISVLYHVGLMQWVILKIAWLMQVTMGTTATETLSVAGNIFVSQTEAPLLIRPYLA
hCNT2   FFGCVVSILYYLGLVQWVVQKVAWFLQITMGTTATETLAVAGNIFVGMTEAPLLIRPYLG
hCNT3   FFSTVMSMLYYLGLMQWIIRKVGWIMLVTTGSSPIESVVASGNIFVGQTESPLLVRPYLP

hCNT1   DMTLSEVHVVMTGGYATIAGSLLGAYISFGIDATSLIAASVMAAPCALALSKLVYPEVEE
hCNT2   DMTLSEIHAVMTGGFATISGTVLGAFIAFGVDASSLISASVMAAPCALASSKLAYPEVEE
hCNT3   YITKSELHAIMTAGFSTIAGSVLGAYISFGVPSSHLLTASVMSAPASLAAAKLFWPETEK

hCNT1   SKFRREEGVKLTYGDAQNLIEAASTGAAISVKVVANIAANLIAFLAVLDFINAALSWLGD
hCNT2   SKFKSEEGVKLPRGKERNVLEAASNGAVDAIGLATNVAANLIAFLAVLAFINAALSWLGE
hCNT3   PKITLKNAMKMESGDSGNLLEAATQGASSSISLVANIAVNLIAFLALLSFMNSALSWFGN

hCNT1   MVDIQGLSFQLICSYILRPVAFLMGVAWEDCPVVAELLGIKLFLNEFVAYQDLSKYKQRR
hCNT2   LVDIQGLTFQVICSYLLRPMVFMMGVEWTDCPMVAEMVGIKFFINEFVAYQQLSQYKNKR
hCNT3   MFDYPQLSFELICSYIFMPFSFMMGVEWQDSFMVARLIGYKTFFNEFVAYEHLSKWIHLR

hCNT1   LAGAEEWVGDRKQWISVRAEVLTTFALCGFANFSSIGIMLGGLTSMVPQRKSDFSQIVLR
hCNT2   LSGMEEWIEGEKQWISVRAEIITTFSLCGFANLSSIGITLGGLTSIVPHRKSDLSKVVVR
hCNT3   KEGGPKFVNGVQQYISIRSEIIATYALCGFANIGSLGIVIGGLTSMAPSRKRDIASGAVR

hCNT1   ALFTGACVSLVNACMAGILYMPRGAEVDCMSLL----NTTLSSSSFEIYQCCREAFQSVN
hCNT2   ALFTGACVSLISACMAGILYVPRGAEADCVSFP----NTSFTNRTYETYMCCRGLFQSTS
hCNT3   ALIAGTVACFMTACIAGILSST-PVDINCHHVLENAFNSTFPGNTTKVIACCQSLLSSTV

hCNT1   ---------------PEFSPEALDNCCRFYNHTICAQ-------
hCNT2   LNGTNPPSFSGPWEDKEFSAMALTNCCGFYNNTVCA--------
hCNT3   AKGPGEVIPG-----GNHSLYSLKGCCTLLNPSTFNCNGISNTF
```

**Figure S3. Evolutionary relationships of NupC with Eukaryota. A.** The amino acid sequence of NupC from *Escherichia coli* (P0AFF2) was subjected to a BLAST search against all Eukaryota in the UniProt KnowledgeBase (http://www.uniprot.org/). Sequences for the top 500 closest proteins were extracted and aligned using the online multiple sequence alignment tool Clustal Omega (http://www.ebi.ac.uk/Tools/msa/clustalo/).[1] The resultant neighbour-joining phylogenetic tree was exported in Newick format and drawn using the online tool iTOL: Interactive Tree of Life (http://itol.embl.de/).[5] **B.** Expansion of the branch containing NupC that also contains two close homologues of NupC, A0A0A2W106 and A0A0A2V7L6, from the soil fungus *Beauveria bassiana*.

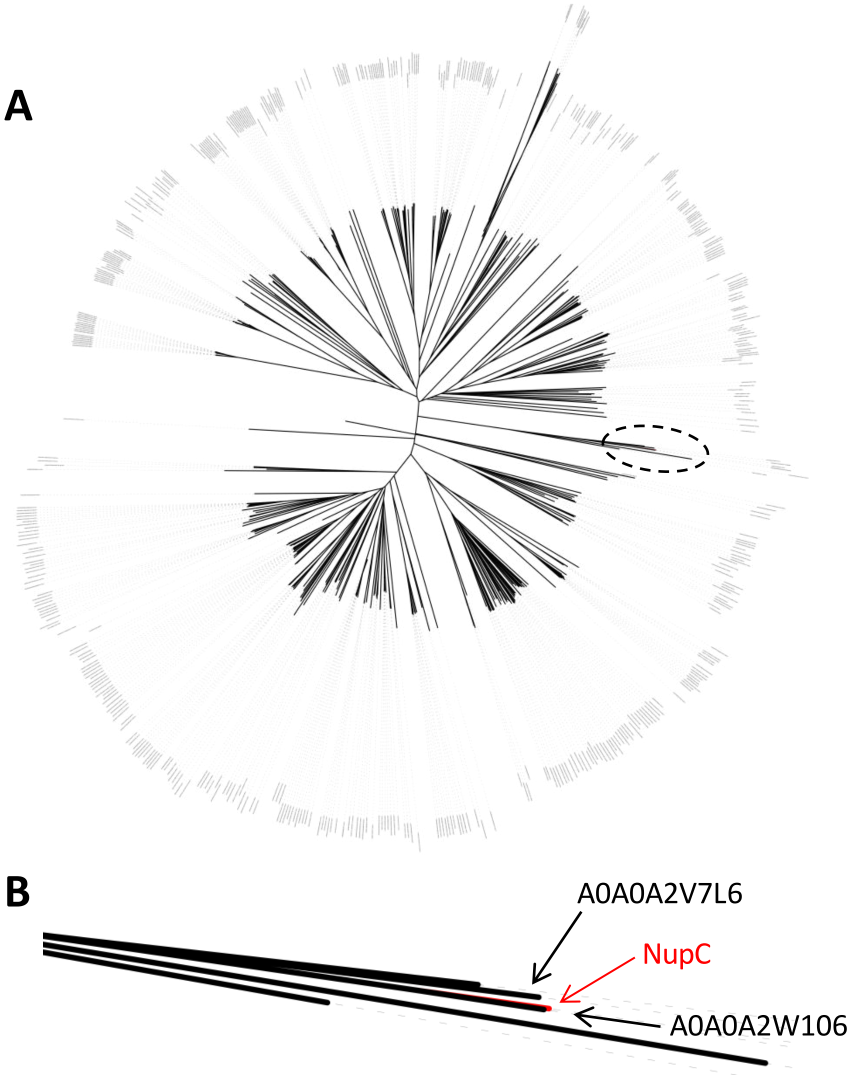

**Figure S4. Conservation of residues in NupC from *Escherichia coli* and vcCNT from *Vibrio cholerae*.** Amino acid sequences of NupC from *Escherichia coli* (P0AFF2) and vcCNT from *Vibrio cholera* (Q9KPL5) were taken from the UniProt KnowledgeBase (http://www.uniprot.org/) and aligned using the online multiple sequence alignment tool Clustal Omega (http://www.ebi.ac.uk/Tools/msa/clustalo/).[1] Residues are coloured to indicate those that are identical (*red*) or highly similar (*blue*). Residues in vcCNT are highlighted to indicate those that contribute to transmembrane spanning helices (TM1-TM8) (*green*), interfacial helices (IH1-IH3) (*cyan*), hairpin re-entrant loops (HP1 and HP2) (*yellow*) and in direct binding interactions with nucleosides (Gly153, Gln154, Thr155, Glu156, Val188, Glu332, Phe366, Asn368 and Ser371) (*pink*) based on crystal structures of vcCNT.[6,7]

```
NupC    ---MDRVLHF--VLALAVVAILALLVSSDRKKIRIRYVIQLLVIEVLLAWFFLNSDVGLG 55
vcCNT   ------MSLFMSCCGMAVLLGIAVLLSSNRKAINLRTVGGAFAIQFSLGAFILYVPWGQE 54

NupC    FVKGFSEMFEKLLGFANEGTNFVFGSMNDQG--------LAFFFLKVLCPIVFISALIGI 107
vcCNT   LLRGFSDAVSNVINYGNDGTSFLFGGLVSGKMFEVFGGGGFIFAFRVLPTLIFFSALISV 114

NupC    LQHIRVLPVIIRAIGFLLSKVNGMGKLESFNAVSSLILGQSENFIAYKDILGKISRNRMY 167
vcCNT   LYYLGVMQWVIRILGGGLQKALGTSRAESMSAAANIFVGQTEAPLVVRPFVPKMTQSELF 174

NupC    TMAATAMSTVSMSIVGAYMTM-LEPKYVVAALVLNMFSTFIVLSLINPYRVD-ASEENIQ 225
vcCNT   AVMCGGLASIAGGVLAGYASMGVKIEYLVAASFMAAPGGLLFAKLMMPETEKPQDNEDIT 234

NupC    MSNL-HEGQSFFEMLGEYILAGFKVAIIVAAMLIGFIALIAALNALFATVTGWFGYS-IS 283
vcCNT   LDGGDDKPANVIDAAAGGASAGLQLALNVGAMLIAFIGLIALINGMLGGIGGWFGMPELK 294

NupC    FQGILGYIFYPIAWVMGVPSSEALQVGSIMATKLVSNEFVAMMDLQKIA-----STLSPR 338
vcCNT   LEMLLGWLFAPLAFLIGVPWNEATVAGEFIGLKTVANEFVAYSQFAPYLTEAAPVVLSEK 354

NupC    AEGIISVFLVSFANFSSIGIIAGAVKGLNEEQGNVVSRFGLKLVYGSTLVSVLSASIAAL 398
vcCNT   TKAIISFALCGFANLSSIAILLGGLGSLAPKRRGDIARMGVKAVIAGTLSNLMAATIAGF 414

NupC    VL-- 400
vcCNT   FLSF 418

Red        Identical residues
Blue       Highly similar residues
TM1-TM8    Transmembrane helices in vcCNT
IH1-IH3    Interfacial helices in vcCNT
HP1-HP2    Hairpin re-entrant loops in vcCNT
Binding    Nucleoside binding in vcCNT
```

**Figure S5. Conservation of residues in nucleoside binding regions of concentrative nucleoside transporters.** Sections representing the structural domains HP1, TM4, HP2 and TM7 in vcCNT taken from a complete amino acid sequence alignment between NupC from *Escherichia coli* (P0AFF2), vcCNT from *Vibrio cholera* (Q9KPL5) and human concentrative nucleoside transporters hCNT1 (O00337), hCNT2 (O43868) and hCNT3 (Q9HAS3). Sequences were taken from the UniProt KnowledgeBase (http://www.uniprot.org/) and aligned using the online multiple sequence alignment tool Clustal Omega (http://www.ebi.ac.uk/Tools/msa/clustalo/).[1] Residues are coloured to indicate those that are identical (*red*) or highly similar (*blue*). Residues in vcCNT are highlighted to indicate those that contribute to the given structural domain (HP1 and HP2) (*yellow*), (TM4 and TM7) (*green*) and in direct binding interactions with nucleosides (Gly153, Gln154, Thr155, Glu156, Val188, Glu332, Phe366, Asn368 and Ser371) (*pink*) based on crystal structures of vcCNT.[6,7]

**HP1**

```
NupC    KVNGMGKLESFNAVSSLILGQSENF 151
vcCNT   KALGTSRAESMSAAANIFVGQTEAP 158
hCNT1   VTMGTTATETLSVAGNIFVSQTEAP 324
hCNT2   ITMGTTATETLAVAGNIFVGMTEAP 318
hCNT3   VTTGSSPIESVVASGNIFVGQTESP 345
```

**TM4**

```
NupC    LGKISRNRMYTMAATAMSTVSMSIV 182
vcCNT   VPKMTQSELFAVMCGGLASIAGGVL 189
hCNT1   LADMTLSEVHVVMTGGYATIAGSLL 355
hCNT2   LGDMTLSEIHAVMTGGFATISGTVL 349
hCNT3   LPYITKSELHAIMTAGFSTIAGSVL 376
```

**HP2**

```
NupC    VPSSEALQVGSIMATKLVSNEFVAM 325
vcCNT   VPWNEATVAGEFIGLKTVANEFVAY 336
hCNT1   VAWEDCPVVAELLGIKLFLNEFVAY 502
hCNT2   VEWTDCPMVAEMVGIKFFINEFVAY 496
hCNT3   VEWQDSFMVARLIGYKTFFNEFVAY 523
```

**TM7**

```
NupC    --STLSPRAEGIISVFLVSFANFSSIGIIA 360
vcCNT   APVVLSEKTKAIISFALCGFANLSSIAILL 376
hCNT1   RKQWISVRAEVLTTFALCGFANFSSIGIML 552
hCNT2   EKQWISVRAEIITTFSLCGFANLSSIGITL 546
hCNT3   VQQYISIRSEIIATYALCGFANIGSLGIVI 573
```

| | |
|---|---|
| **Red** | Identical residues |
| **Blue** | Highly similar residues |
| HP1+HP2 | Hairpin re-entrant loops in vcCNT |
| TM4+TM7 | Transmembrane helices in vcCNT |
| Binding | Nucleoside binding in vcCNT |

**Figure S6. Protein sequence alignment between concentrative nucleoside transporters.** Amino acid sequences of NupC from *Escherichia coli* (P0AFF2), vcCNT from *Vibrio cholera* (Q9KPL5) and human concentrative nucleoside transporters hCNT1 (O00337), hCNT2 (O43868) and hCNT3 (Q9HAS3) were taken from the UniProt KnowledgeBase (http://www.uniprot.org/) and aligned using the online multiple sequence alignment tool Clustal Omega (http://www.ebi.ac.uk/Tools/msa/clustalo/).[1] Residues are coloured to indicate those that are identical (*red*) and highly similar (*blue*). vcCNT residues are highlighted to show those that contribute to transmembrane spanning helices (TM1-TM8, *green*), interfacial helices (IH1-IH3, *cyan*) and hairpin re-entrant loops (HP1 and HP2, *yellow*) based on crystal structures of vcCNT.[6,7]

```
NupC   ------------------------------------------------------------
vcCNT  ------------------------------------------------------------
hCNT1  -------------------------MENDPSRRRESISLTPVAKGLENMGADFLESLEEG 35
hCNT2  ---------------------------MEKASGRQSIALSTVETGTVNPGLELMEKEVEP 33
hCNT3  MELRSTAAPRAEGYSNVGFQNEENFLENENTSGNNSIRSRAVQSREHTNTKQDEEQVTVE 60

NupC   ------------------------------------------------------------
vcCNT  ------------------------------------------------------------
hCNT1  Q-LPRSDLSPAEIRSSWSEAAPKPFSRWRNLQPALRARSFCREHMQLFRWIGTGLLCTGL 94
hCNT2  EGSKRTDAQGHSLGDG---LGPSTY-QRRSRWPFSKARSFCKTHASLFKKILLGLLCLAY 89
hCNT3  QDSPRNREH---MEDDDEEMQQKG----CLERRYDTVCGFCRKHKTTLRHIIWGILLAGY 113

NupC   ------------------------------------------------------------
vcCNT  ------------------------------------------------------------
hCNT1  SAFLLVACLLDFQRALALFVLTCVVLTFLGHRLLKRLLGPKLRRFLKPQGH--PRLLLWF 153
hCNT2  AAYLLAACILNFQRALALFVITCLVIFVLVHSFLKKLLGKKLTRCLKPFEN--SRLRLWT 147
hCNT3  LVMVISACVLNFHRALPLFVITVAAIFFVVWDHLMAKYEHRIDEMLSPGRRLLNSHWFWL 173

NupC   ------------------------MDRVLHFVLALAVVAILALLVSSDRKKIRIRYVIQL 36
vcCNT  ------------------GPAVPRMSLFMS-CCGMAVLLGIAVLLSSNRKAINLRTVGGA 41
hCNT1  KRGLALAAFLGLVLWLSLDTSQR-PEQLVS-FAGICVFVALLFACSKHHCAVSWRAVSWG 211
hCNT2  KWVFAGVSLVGLILWLALDTAQR-PEQLIP-FAGICMFILILFACSKHHSAVSWRTVFSG 205
hCNT3  KWVIWSSLVLAVIFWLAFDTAKLGQQQLVS-FGGLIMYIVLLFLFSKYPTRVYWRPVLWG 232

NupC   LVIEVLLAWFFLNSDVGLGFVKGFSEMFEKLLGFANEGTNFVFGSMNDQ--------GLA 88
vcCNT  FAIQFSLGAFILYVPWGQELLRGFSDAVSNVINYGNDGTSFLFGGLVSGKMFEVFGGGGF 101
hCNT1  LGLQFVLGLLVIRTEPGFIAFEWLGEQIRIFLSYTKAGSSFVFGEALVK----------D 261
hCNT2  LGLQFVFGILVIRTDLGYTVFQWLGEQVQIFLNYTVAGSSFVFGDTLVK----------D 255
hCNT3  IGLQFLLGLLILRTDPGFIAFDWLGRQVQTFLEYTDAGASFVFGEKYKD----------H 282

NupC   FFFLKVLCPIVFISALIGILQHIRVLPVIIRAIGFLLSKVNGMGKLESFNAVSSLILGQS 148
vcCNT  IFAFRVLPTLIFFSALISVLYYLGVMQWVIRILGGGLQKALGTSRAESMSAAANIFVGQT 161
hCNT1  VFAFQVLPIIVFFSCVISVLYHVGLMQWVILKIAWLMQVTMGTTATETLSVAGNIFVSQT 321
hCNT2  VFAFQALPIIIFFGCVVSILYYLGLVQWVVQKVAWFLQITMGTTATETLAVAGNIFVGMT 315
hCNT3  FFAFKVLPIVVFFSTVMSMLYYLGLMQWIIRKVGWIMLVTTGSSPIESVVASGNIFVGQT 342

NupC   ENFIAYKDILGKISRNRMYTMAATAMSTVSMSIVGAYMTMLEP-KYVVAALVLNMFSTFI 207
vcCNT  EAPLVVRPFVPKMTQSELFAVMCGGLASIAGGVLAGYASMGVKIEYLVAASFMAAPGGLL 221
hCNT1  EAPLLIRPYLADMTLSEVHVVMTGGYATIAGSLLGAYISFGIDATSLIAASVMAAPCALA 381
hCNT2  EAPLLIRPYLGDMTLSEIHAVMTGGFATISGTVLGAFIAFGVDASSLISASVMAAPCALA 375
hCNT3  ESPLLVRPYLPYITKSELHAIMTAGFSTIAGSVLGAYISFGVPSSHLLTASVMSAPASLA 402

NupC   VLSLLINPYRVD---ASEENIQMSNLHEGQSFFEMLGEYILAGFKVAIIVAAMLIGFIALI 264
vcCNT  FAKLMMPETEKPQDNEDITLDGG-DDKPANVIDAAAGGASAGLQLALNVGAMLIAFIGLI 280
hCNT1  LSKLVYPEVEESKFRREEGVKLT-YGDAQNLIEAASTGAAISVKVVANIAANLIAFLAVL 440
hCNT2  SSKLAYPEVEESKFKSEEGVKLP-RGKERNVLEAASNGAVDAIGLATNVAANLIAFLAVL 434
hCNT3  AAKLFWPETEKPKITLKNAMKME-SGDSGNLLEAATQGASSSISLVANIAVNLIAFLALL 461
```

```
NupC    AALNALFATVTGWFGYS-ISFQGILGYIFYPIAWVMGVPSSEALQVGSIMATKLVSNEFV 323
vcCNT   ALINGMLGGIGGWFGMPELKLEMLLGWLFAPLAFLIGVPWNEATVAGEFIGLKTVANEFV 340
hCNT1   DFINAALSWLGDMVDIQGLSFQLICSYILRPVAFLMGVAWEDCPVVAELLGIKLFLNEFV 500
hCNT2   AFINAALSWLGELVDIQGLTFQVICSYLLRPMVFMMGVEWTDCPMVAEMVGIKFFINEFV 494
hCNT3   SFMNSALSWFGNMFDYPQLSFELICSYIFMPFSFMMGVEWQDSFMVARLIGYKTFFNEFV 521

NupC    AMMDLQKIA---------------STLSPRAEGIISVFLVSFANFSSIGIIAGAVKGLNE 368
vcCNT   AYSQFAPYLTEA----------APVVLSEKTKAIISFALCGFANLSSIAILLGGLGSLAP 390
hCNT1   AYQDLSKYKQRRLAGAEEWVGDRKQWISVRAEVLTTFALCGFANFSSIGIMLGGLTSMVP 560
hCNT2   AYQQLSQYKNKRLSGMEEWIEGEKQWISVRAEIITTFSLCGFANLSSIGITLGGLTSIVP 554
hCNT3   AYEHLSKWIHLRKEGGPKFVNGVQQYISIRSEIIATYALCGFANIGSLGIVIGGLTSMAP 581

NupC    EQGNVVSRFGLKLVYGSTLVSVLSASIAALVL---------------------------- 400
vcCNT   KRRGDIARMGVKAVIAGTLSNLMAATIAGFFLSF-------------------------- 424
hCNT1   QRKSDFSQIVLRALFTGACVSLVNACMAGILYMPRGAEVDCMSLL----NTTLSSSSFEI 616
hCNT2   HRKSDLSKVVVRALFTGACVSLISACMAGILYVPRGAEADCVSFP----NTSFTNRTYET 610
hCNT3   SRKRDIASGAVRALIAGTVACFMTACIAGILSST-PVDINCHHVLENAFNSTFPGNTTKV 640

NupC    --------------------------------------------------------
vcCNT   --------------------------------------------------------
hCNT1   YQCCREAFQSVN---------------PEFSPEALDNCCRFYNHTICAQ------- 650
hCNT2   YMCCRGLFQSTSLNGTNPPSFSGPWEDKEFSAMALTNCCGFYNNTVCA-------- 658
hCNT3   IACCQSLLSSTVAKGPGEVIPG-----GNHSLYSLKGCCTLLNPSTFNCNGISNTF 691
```

**Figure S7. Membrane topology predictions for NupC.** The amino acid sequence of NupC from *Escherichia coli* (P0AFF2) was taken from the UniProt KnowledgeBase (http://www.uniprot.org/) and analysed by the membrane topology prediction tools TOPCONS (http://topcons.cbr.su.se/pred/)[8] (**A**), TMHMM (http://www.cbs.dtu.dk/services/TMHMM/)[9] (**B**) and CCTOP (http://cctop.enzim.ttk.mta.hu/)[10] (**C**).

#### A. TOPCONS

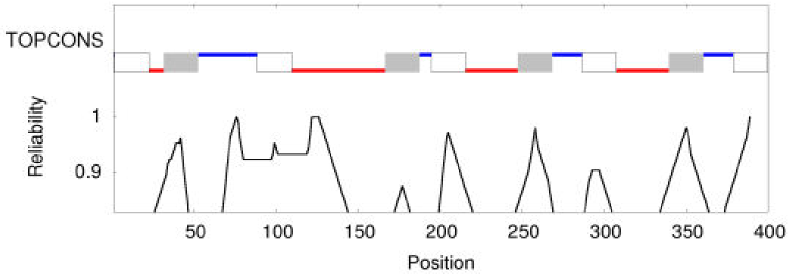


TM helices (white/grey boxes)
Extracellular regions (blue)
Intracellular regions (red)

#### B. TMHMM

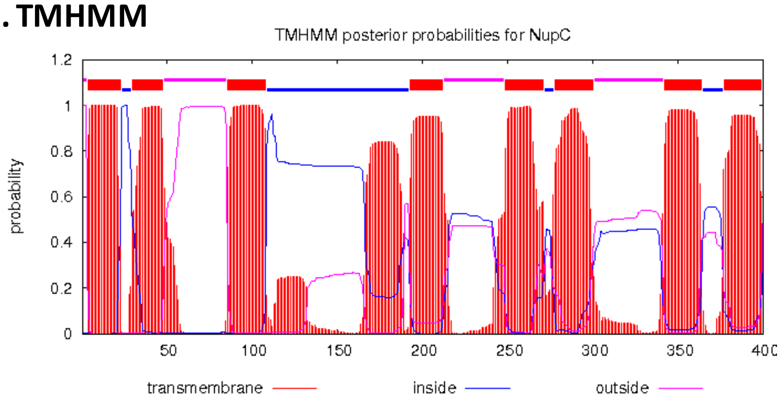


TM helices (red boxes)
Extracellular regions (pink)
Intracellular regions (blue)

#### C. CCTOP

<Topology numTM="9" reliability="78.0708">
<Region from="1" to="3" loc="I"/>
<Region from="4" to="22" loc="M"/>
<Region from="23" to="31" loc="O"/>
<Region from="32" to="50" loc="M"/>
<Region from="51" to="86" loc="I"/>
<Region from="87" to="108" loc="M"/>
<Region from="109" to="167" loc="O"/>
<Region from="168" to="189" loc="M"/>
<Region from="190" to="193" loc="I"/>
<Region from="194" to="212" loc="M"/>
<Region from="213" to="248" loc="O"/>
<Region from="249" to="270" loc="M"/>
<Region from="271" to="277" loc="I"/>
<Region from="278" to="299" loc="M"/>
<Region from="300" to="340" loc="O"/>
<Region from="341" to="362" loc="M"/>
<Region from="363" to="377" loc="I"/>
<Region from="378" to="399" loc="M"/>
<Region from="400" to="400" loc="O"/>

TM helices (M, red)
Extracellular regions (O)
Intracellular regions (I)

**Figure S8. Predictions of transmembrane helices in NupC from *Escherichia coli* and in vcCNT from *Vibrio cholerae*.** Amino acid sequences of NupC from *Escherichia coli* (P0AFF2) (*left*) and vcCNT from *Vibrio cholera* (Q9KPL5) (*right*) were taken from the UniProt KnowledgeBase (http://www.uniprot.org/) and analysed by the membrane topology prediction tools TOPCONS (http://topcons.cbr.su.se/pred/),[8] TMHMM (http://www.cbs.dtu.dk/services/TMHMM/)[9] and CCTOP (http://cctop.enzim.ttk.mta.hu/).[10] Coloured highlighting is used to show the positions of predicted full transmembrane spanning helices (*grey*), the positions of transmembrane helices based on the crystal structure of vcCNT (Figure 3) (*green*) and the positions of positively charged residues at physiological pH (arginine and lysine) (*red*).

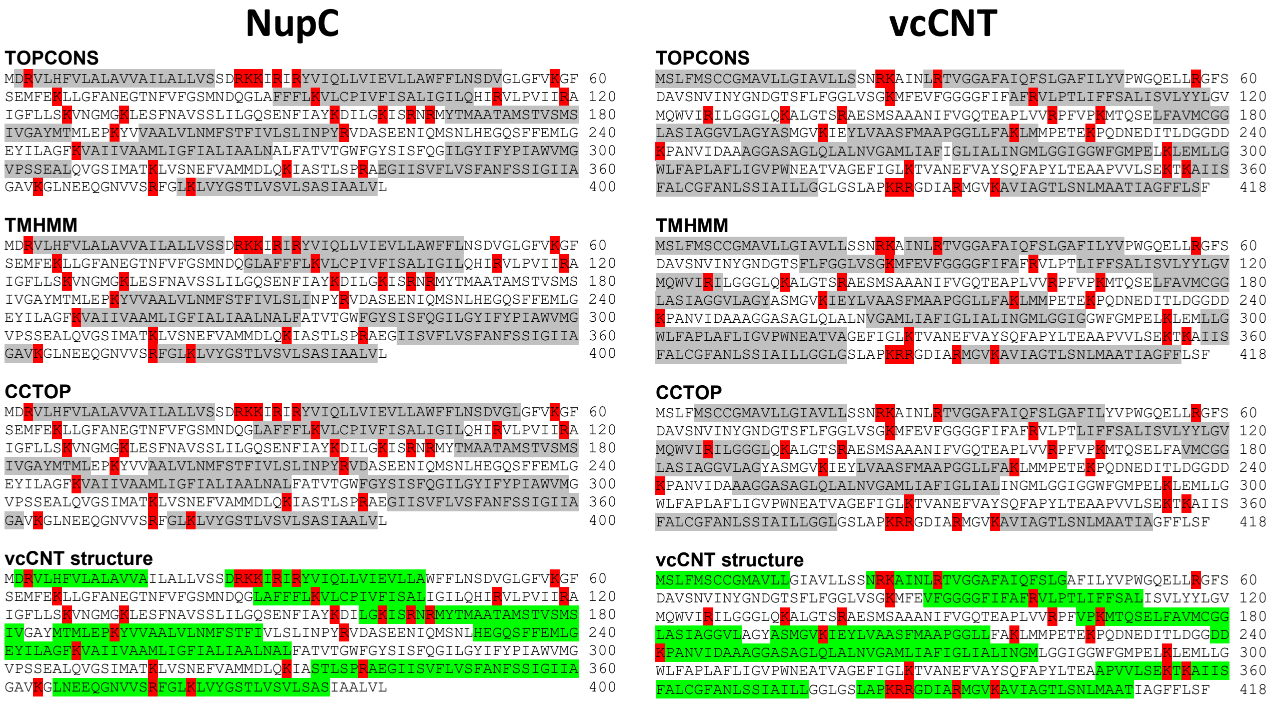

**Figure S9. Membrane topology predictions for vcCNT.** The amino acid sequence of vcCNT from *Vibrio cholera* (Q9KPL5) was taken from the UniProt KnowledgeBase (http://www.uniprot.org/) and analysed by the membrane topology prediction tools TOPCONS (http://topcons.cbr.su.se/pred/)[8] (**A**), TMHMM (http://www.cbs.dtu.dk/services/TMHMM/)[9] (**B**) and CCTOP (http://cctop.enzim.ttk.mta.hu/)[10] (**C**).

#### A. TOPCONS

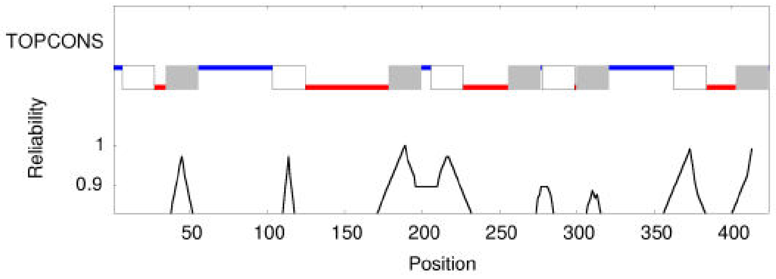


TM helices (white/grey boxes)
Extracellular regions (blue)
Intracellular regions (red)

#### B. TMHMM

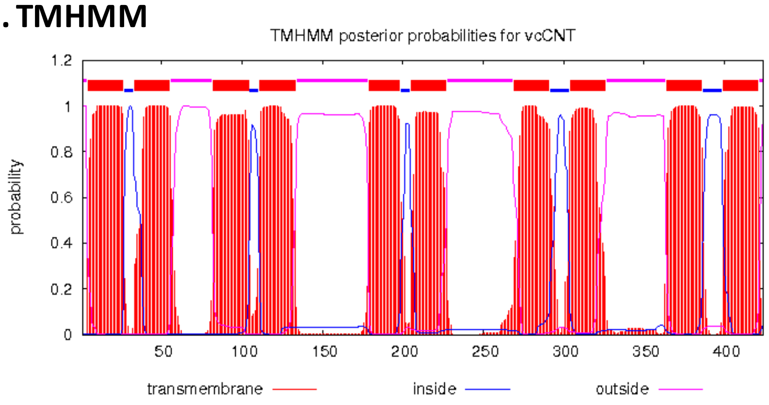


TM helices (red boxes)
Extracellular regions (pink)
Intracellular regions (blue)

#### C. CCTOP

```
<Topology numTM="8" reliability="84.7869">
<Region from="1" to="10" loc="O"/>
<Region from="11" to="26" loc="M"/>
<Region from="27" to="36" loc="I"/>
<Region from="37" to="53" loc="M"/>
<Region from="54" to="110" loc="O"/>
<Region from="111" to="138" loc="M"/>
<Region from="139" to="148" loc="I"/>
<Region from="149" to="165" loc="L"/>
<Region from="166" to="181" loc="I"/>
<Region from="182" to="197" loc="M"/>
<Region from="198" to="207" loc="O"/>
<Region from="208" to="223" loc="M"/>
<Region from="224" to="254" loc="I"/>
<Region from="255" to="282" loc="M"/>
<Region from="283" to="304" loc="O"/>
<Region from="305" to="317" loc="L"/>
<Region from="318" to="329" loc="O"/>
<Region from="330" to="342" loc="L"/>
<Region from="343" to="366" loc="O"/>
<Region from="367" to="385" loc="M"/>
<Region from="386" to="402" loc="I"/>
<Region from="403" to="418" loc="M"/>
<Region from="419" to="424" loc="O"/>
```

TM helices (M, red)
Extracellular regions (O)
Intracellular regions (I)

**Figure S10. Amplified expression of wild-type NupC and mutants in inner membrane preparations.** Coomassie blue-stained SDS-PAGE separation of proteins (30 μg) in inner membrane preparations from *E. coli* BL21(DE3) cells harbouring plasmid pGJL16 for expressing WT NupC or constructs for expressing mutants S142C, G146C and E149C following growth in minimal medium and induction with IPTG. The mobilities of molecular weight (MW) marker proteins are shown and the arrow indicates the position of NupC.

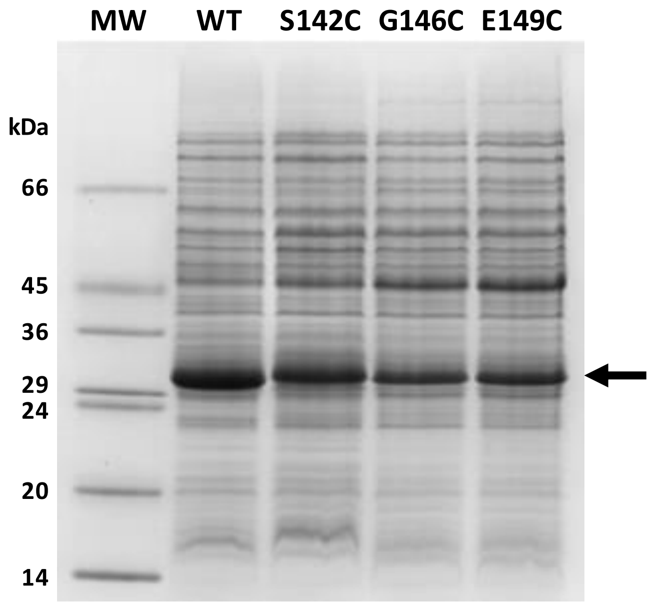

**Figure S11. Initial rates of NupC-mediated [$^{14}$C]uridine uptake into energised *E. coli* cells.** Uptake of [$^{14}$C]uridine (5 μM and 100 μM) after time points of 5, 10, 15 and 20 seconds into transport energised *E. coli* BL21(DE3) cells harbouring plasmid pGJL16 for expressing WT NupC from a culture grown in minimal medium and induced with IPTG. Data points represent the average of duplicate measurements.

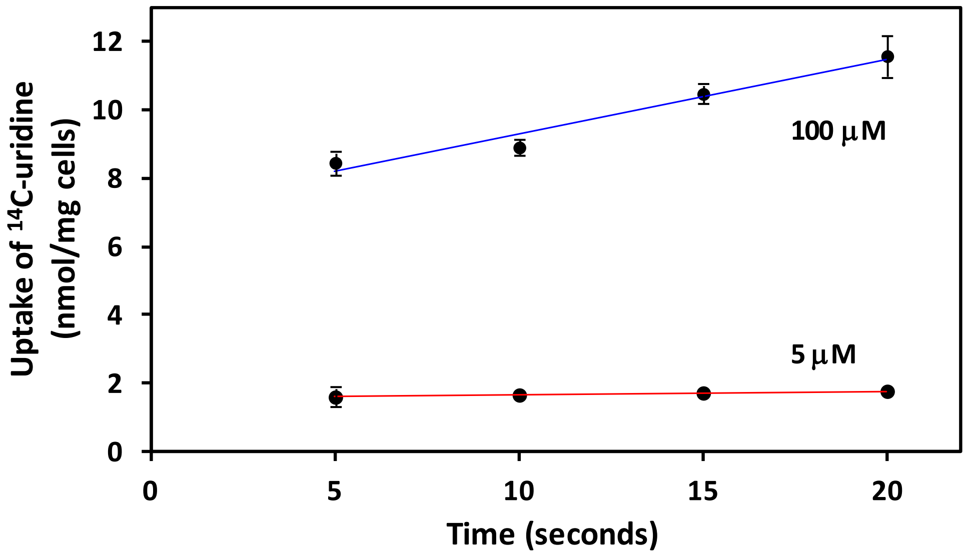

**Figure S12. NupC-mediated uptake of [$^{14}$C]adenosine and [$^{14}$C]uridine into energised *E. coli* cells.** Uptake of [$^{14}$C]adenosine (50 μM) and [$^{14}$C]uridine (50 μM) after 15 seconds and 2 minutes into transport energised *E. coli* BL21(DE3) cells harbouring empty plasmid pTTQ18 (no transporter gene insert) induced with IPTG (0.05 mM) (**A**) or plasmid pGJL16 for expressing WT NupC without induction (**B**) or induced with IPTG (**C**) from cultures grown in minimal medium. Data points represent the average of duplicate measurements.

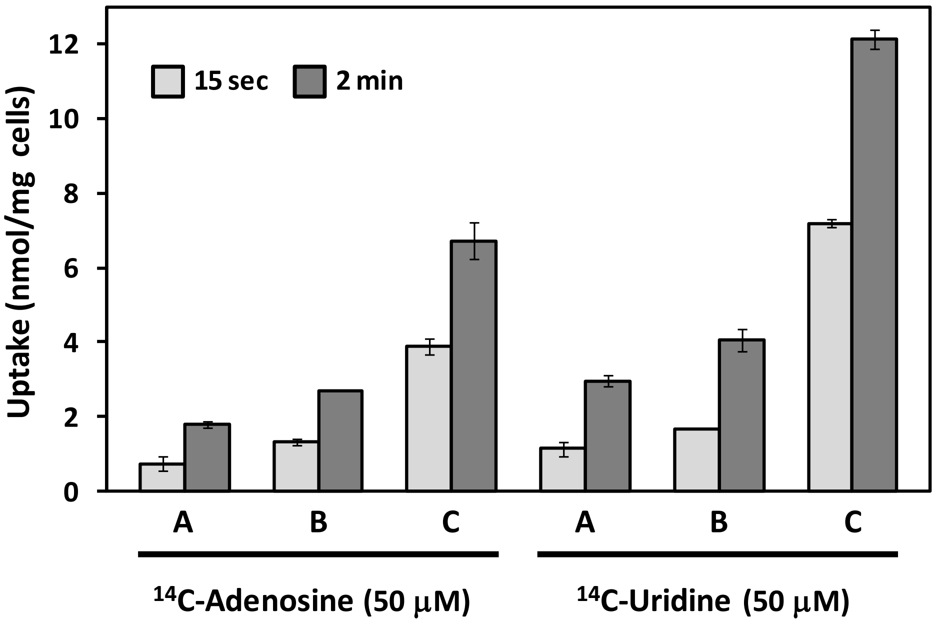
